# Sensory Feedback and Central Neuronal Interactions in Mouse Locomotion

**DOI:** 10.1101/2023.10.31.564886

**Authors:** Yaroslav I. Molkov, Guoning Yu, Jessica Ausborn, Julien Bouvier, Simon M. Danner, Ilya A. Rybak

**Author notes:** Corresponding Authors: Yaroslav Molkov, PhD, Department of Mathematics and Statistics, Neuroscience Institute, Georgia State University, 25 Park Place, Atlanta, GA 30303, USA, Ilya A. Rybak, PhD, Department of Neurobiology and Anatomy, Drexel University College of Medicine, 2900 Queen Lane, Philadelphia, PA 19129, USA.

## Abstract

Locomotion is a complex process involving specific interactions between the central neural controller and the mechanical components of the system. The basic rhythmic activity generated by locomotor circuits in the spinal cord defines rhythmic limb movements and their central coordination. The operation of these circuits is modulated by sensory feedback from the limbs providing information about the state of the limbs and the body. However, the specific role and contribution of central interactions and sensory feedback in the control of locomotor gait and posture remain poorly understood. We use biomechanical data on quadrupedal locomotion in mice and recent findings on the organization of neural interactions within the spinal locomotor circuitry to create and analyze a tractable mathematical model of mouse locomotion. The model includes a simplified mechanical model of the mouse body with four limbs and a central controller composed of four rhythm generators, each operating as a state machine controlling the state of one limb. Feedback signals characterize the load and extension of each limb as well as postural stability (balance). We systematically investigate and compare several model versions and compare their behavior to existing experimental data on mouse locomotion. Our results highlight the specific roles of sensory feedback and some central propriospinal interactions between circuits controlling fore and hind limbs for speed-dependent gait expression. Our models suggest that postural imbalance feedback may be critically involved in the control of swing-to-stance transitions in each limb and the stabilization of walking direction.

## 1 Introduction

Locomotion in quadrupeds is a complex process that involves the coordination of movements of four limbs actuated by numerous muscles. This coordination defines locomotor gaits and ensures postural stability during locomotion at the desired velocity and direction, as well as their changes. The basic pattern of limb movements during locomotion is produced by neural circuitry in the spinal cord, commonly referred to as a Central Pattern Generator (CPG) [1-5]. The locomotor CPG represents a central neural controller comprising rhythm-generating and limb-coordinating circuits [2, 3, 5-7]. CPG operation is modulated by supraspinal inputs and sensory feedback that informs the central controller about the state of the limbs and the body (posture) [8-11]. Interlimb coordination is provided, at least in part, by left-right and fore-hind interactions between the rhythm-generating circuits controlling each limb. Such central coordination of left-right and fore-hind motor activity is present during so-called *fictive locomotion*, the locomotor-like activity generated in the absence of limb movements and motion-dependent feedback from the limbs [12-16]. However, during actual locomotion, sensory feedback from the limbs, which reflects the state of limb muscles, limb/body mechanics, and interactions with the environment, can modulate or even override the locomotor oscillations generated by the spinal circuits and their coordination. The specific interactions between the central controller and sensory feedback, as well as the role of different feedback types in regulating locomotor speed, gait, postural stability, and movement direction under different experimental conditions, remain contradictory and poorly understood [17-23]. Moreover, some theories almost exclusively rely on the critical role of sensory feedback in the timing of locomotor phase transitions and interlimb coordination [24-27], hence devaluing the role of central mechanisms. In addition (and importantly), most previous modeling studies considered and analyzed the role of feedback for locomotion on a step-by-step basis without considering its role in the maintenance of the direction of movement. In these models, the direction of movement was explicitly or implicitly restricted to a straight-line with the body aligned with the direction of movement. These limitations prevented exploring the ability of the system to maintain a constant direction of movement and the roles and effects of body displacement and rotation.

In this study, we address the above issues using a tractable mathematical model describing body movements and locomotion in a two-dimensional space. The model combines key biomechanical data from studies of quadrupedal locomotion in mice with recent data on neural interactions within the spinal locomotor circuitry. The central neural controller (or the CPG) is represented by a minimal model comprising four rhythm generators (RGs), each controlling a single limb. The four RGs receive a minimal set of feedback signals resulting from the loading and extension of each limb. These signals play a critical role in controlling the locomotor gait and contribute to maintaining postural stability during locomotion. We comparatively investigate several model versions by analyzing their ability to locomote with different speeds and phase durations and to maintain the direction of movement and by comparing their behavior with existing experimental data on mouse locomotion [28-31].

Based on our modeling, we suggest that in freely moving mice, the stance-to-swing transition in each limb is directly triggered by feedback to the RG from the homonymous limb (the limb controlled by the same RG), while the swing-to-stance transition in each limb may require a common posture/balance-dependent feedback signal. We introduce and consider such a feedback signal based on the balance of the body. This feedback signal is received by all RGs and depending on each limb’s state, may induce a touchdown of the swinging limb and thus control its swing duration. We show that this feedback provides stability of the walking direction, an important feature that has not been explicitly considered in previous studies. We also show that the proposed mechanisms for phase transitions, together with some central interactions within the spinal locomotor circuitry, define the locomotor gait and its dependence on the locomotor speed.

Although the model is primarily based on data and characteristics from mice, it can be scaled for other quadrupedal animals and be used as a convenient theoretical framework that couples neural dynamics with mathematically tractable biomechanics.

## 2 Model description

### 2.1 Model of the CPG

We implement the CPG as a set of four rhythm generators (RGs), each controlling one limb. Each RG operates in a state machine regime, so that at every given moment in time it can be in one of two states: swing or stance (Fig. 1). During stance, the end-effector of the limb (paw), which is controlled by the corresponding RG, is assumed to be on the ground hence providing support for the body. During swing, the limbs instantaneously move to their target position relative to the body and await touchdown. Transitions between the states can occur autonomously due to central (spinal) interactions between the RGs or due to somatosensory feedback. As described below, we consider two fundamental mechanisms of stance-to-swing transitions (liftoff), limb extension and its unloading, combined with different mechanisms of swing-to-stance transitions.

**Figure 1.**
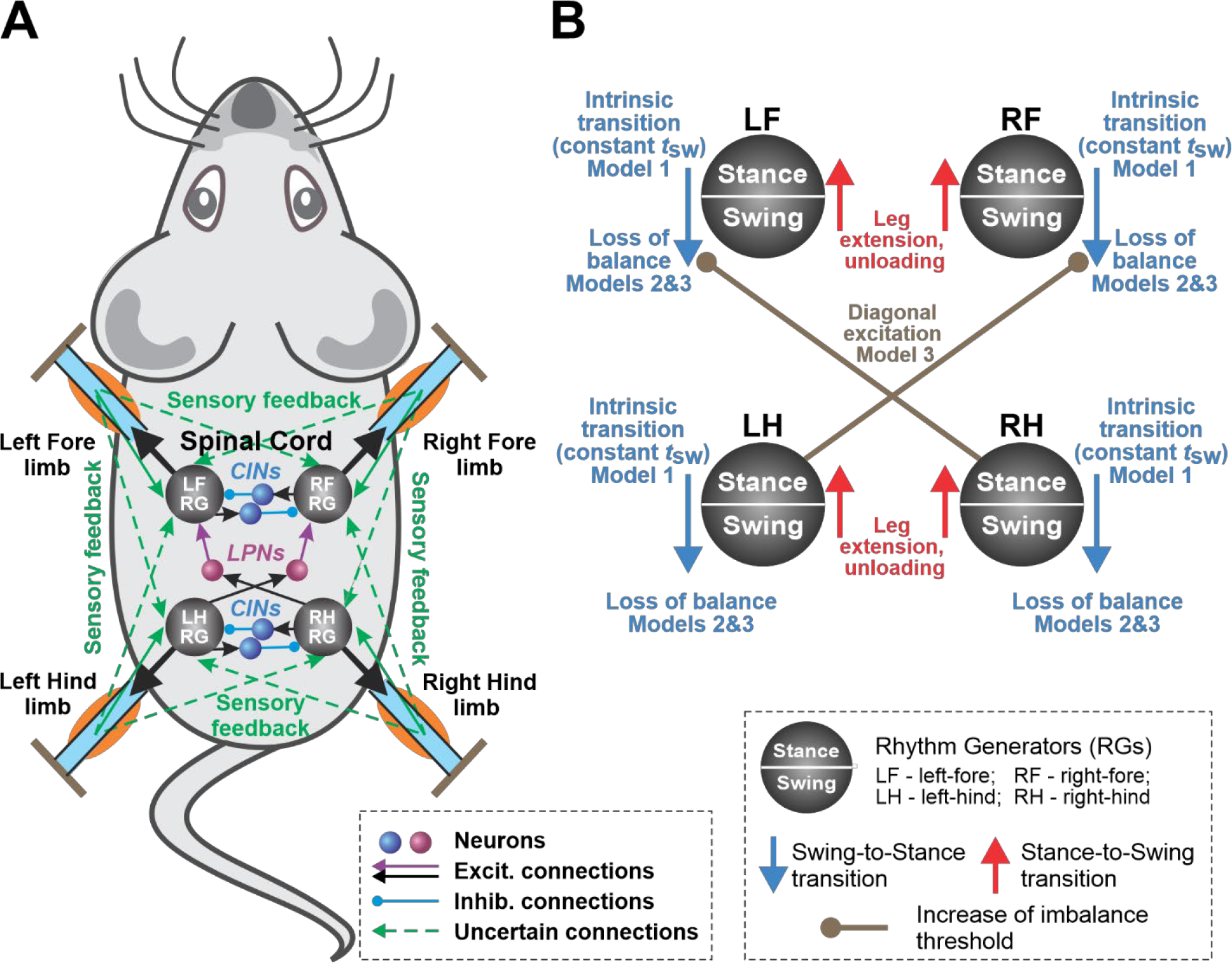
“Central Neural controller” (CPG). **A:** The CPG is composed of four rhythm generators (RGs), each controlling one limb. Central interactions between the RGs are provided by left-right commissural interneurons (CINs) and fore-hind long propriospinal neurons (LPNs). **B:** Each RG is modeled as a ‘*state machine*’ that can be in one of two states: “Swing” or “Stance”, which define the state of the limb controlled by the corresponding RG. Transitions between the states can be defined intrinsically (“Intrinsic transition”) and/or depend on central interactions between the RGs as well as on sensory signals characterizing the state of the corresponding leg (“Leg extension, unloading”) and the body balance (“Loss of balance”).

### 2.2 Model of the body and limbs

For the mouse’s body, we define a rigid frame with length *L* and width 2*h* (Fig. 2). The plane of this frame, also referred to as the *horizontal plane*, is parallel to the ground, and therefore the normal direction to this plane is the *vertical direction*. In terms of mass distribution, the body is considered as a uniform rigid rod of length *L* with the center of mass (COM) at the midpoint of the body as shown in Fig. 2. The distance from the horizontal plane to the ground is defined to be the height *H* of the mouse’s COM.

**Figure 2.**
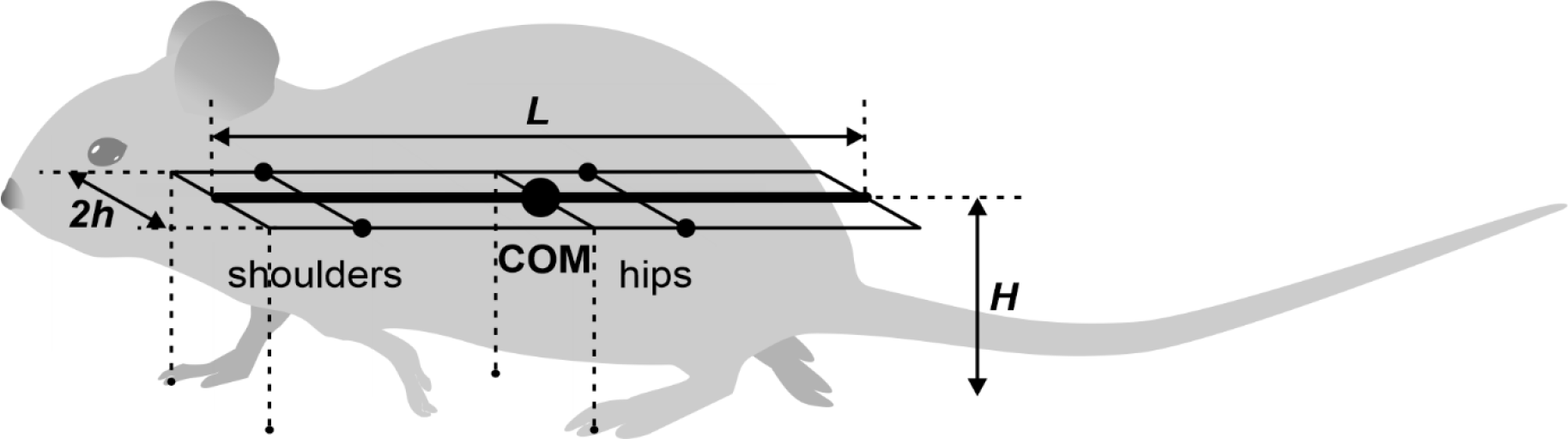
The mouse body is approximated by a rigid rod with a length of *L* with uniform linear density. The left and right “shoulder” and “hip” joints are located in the horizontal plane, each at the same distance *h* from the rod forming the body frame *L* × 2*h*. The initial positions of the limbs during stance (relative to the body) are at the ground projections of the front and middle (crossing the COM) frame segments for the fore and hind limbs, respectively (see the tips of the dashed lines on the ground). The same positions are also considered to be the corresponding target positions of the paws during swing.

The body weight is supported by the limbs that are in stance. The initial positions of the paws in stance correspond to the ground projections of the front of the body’s frame for the forelimbs and of its center (crossing the COM) for the hind limbs (Fig. 2). The initial stance positions of the left and right limbs are displaced from the body’s centerline by distance *h* to the left and to the right. These initial points serve as target positions in the horizontal plane for the corresponding paws during swing. The maximal possible limb displacement in the horizontal plane from its initial position in the coordinate system associated with the body (maximal limb displacement) is equal to *D*.

### 2.3 Equations of motion

Let *m* be the mass of the mouse and *g* be the gravitational acceleration. The external forces considered are the gravity force, friction force and the ground reaction forces. We decompose the forces (where applicable) into components parallel (horizontal) and perpendicular (vertical) to the horizontal plane. Unless otherwise stated, we indicate vectors by bold letters and their magnitudes using the same notations in a regular font.

#### 2.3.1 Horizontal forces and yaw torques

Let ***F***_*i*_ denote the component of ground reaction forces in the horizontal plane (2D vector) for limb *i* (*i* = 1 for the left fore (LF), 2 for the right fore (RF), 3 for the left hind (LH), and 4 for the right hind (RH) limbs). We assume that every paw touching the ground creates a propulsion force with magnitude *F*_0_ (same for all limbs on the ground) directed along the body in the rostral direction. *F*_0_ is an important parameter of the model affecting the locomotor speed. Hereinafter, we refer to this parameter as *propulsion force*.

During locomotion, energy losses are associated with various factors many of which are not explicitly represented in our simplified model. To account for energy dissipation, we introduce an equivalent viscous friction force that linearly depends on the velocity, i.e., ***F***_*f*_ = −*λ****v***_*c*_, where ***v***_*c*_ is the COM velocity vector in the horizontal plane and *λ* is the coefficient of kinetic friction (see the table of parameter values).

When only two limbs support the body, as shown in Fig. 3, the COM movement in the horizontal plane is affected by gravity. We connect the positions of these two limbs by a *supporting line segment*. Let ***z*** be the distance vector from the supporting line segment to the COM projection on the ground and let *φ* be the angle of inclination (Fig. 3). The model can be seen as an inverted pendulum pivoted at the supporting line segment governed by the following equation:

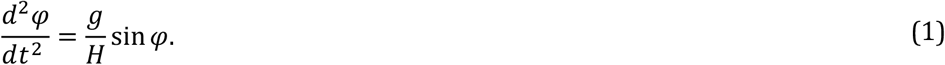

After considering that sin *φ* = *z*/*H* and linearizing Eq. (1):

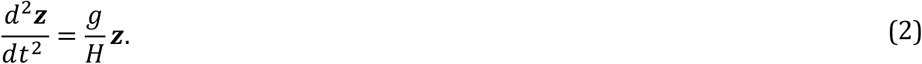

This acceleration multiplied by the mass *m* represents an *inverted pendulum force*, which we denote here by ***F***_*p*_. It can be expressed in terms of the distance vector ***z*** as follows.

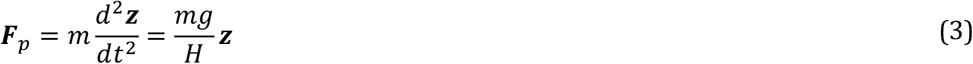

If more than two limbs are on the ground, we assume that the COM is always inside of the supporting triangle or quadrangle, and ***F***_*p*_ = **0**.

**Figure 3.**
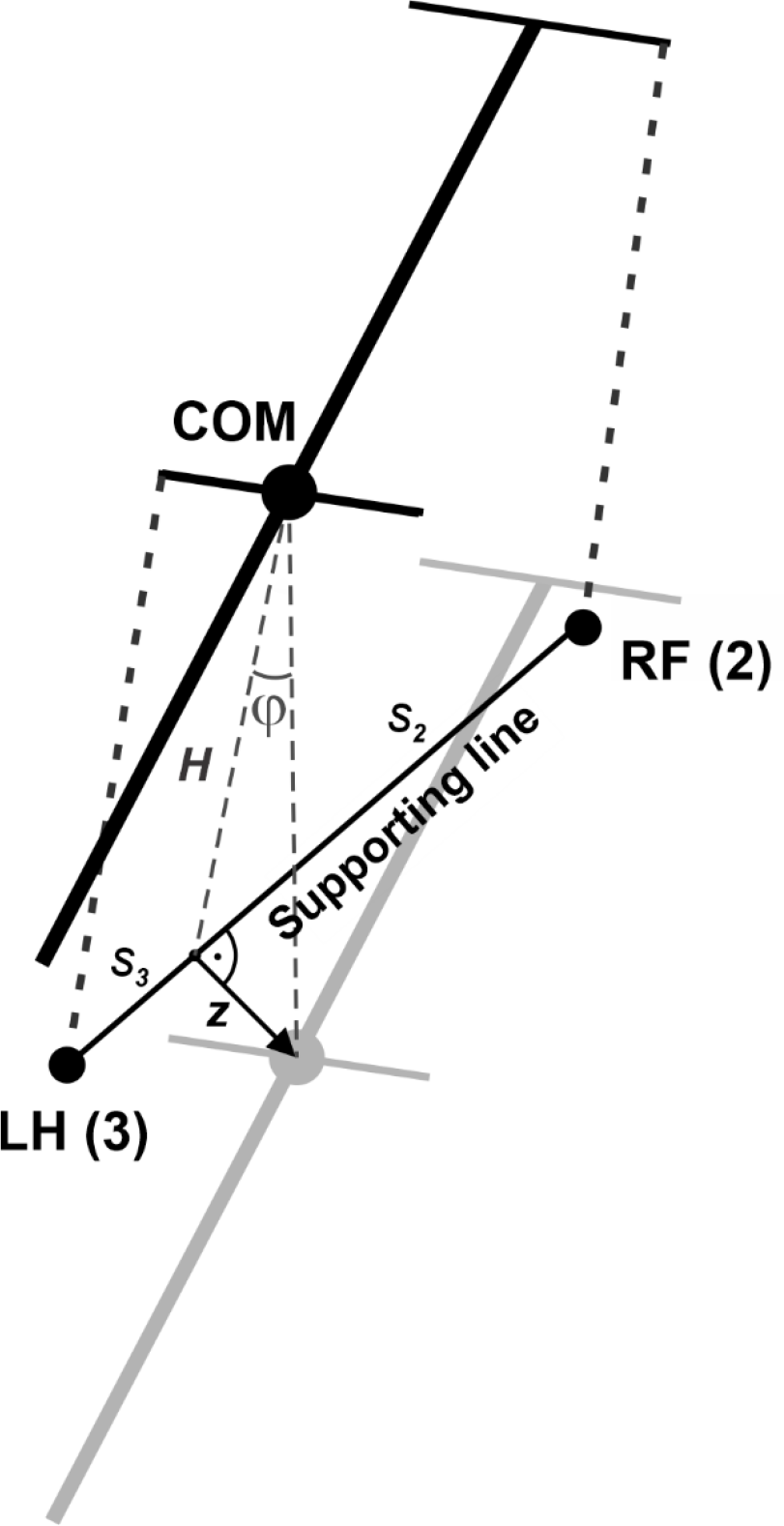
Inverted pendulum dynamics during two-leg support. During two-leg support phases, the body weight cannot be fully supported by the limbs on the ground. In this case, we describe the effect of the gravitational force on the horizontal center of mass (COM) movement by the linearized inverted pendulum model. In the example shown, the right fore (RF (2)) and left hind (LH (3)) limbs are on the ground providing support, while the left fore and right hind limbs (not shown) are in swing. Since the COM is displaced from the supporting line (***z*** is the COM displacement vector in the horizontal plane, *φ* is the corresponding angle between the projection of the COM onto the line of support and the vertical), the gravitational force creates a rolling torque about this line. We approximate this torque as an equivalent horizontal force pushing the COM in the direction perpendicular to the supporting line (along ***z***). See text for details.

By Second Newton’s Law, the velocity *v*_*c*_ obeys the following equation:

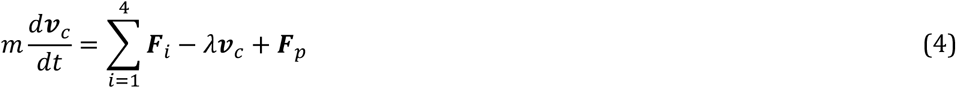

Since the propulsion forces are displaced from the axis of the body, they will contribute to the yaw torque leading to the rotation of the body around the COM in a horizontal plane. The moment of inertia *I* of a uniform rod of length *L* about the COM is

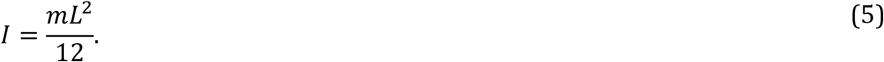

The gravity force is applied at the COM, thus creating zero torque. Let ***r***_*c*_ be the position vector of the COM in the horizontal plane, and ***r***_*i*_ be the position vector of paw *i* on the ground. Then, the distance vector from the COM to paw *i* is

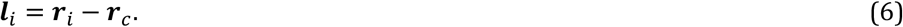

The yaw torque that each propulsion force creates is

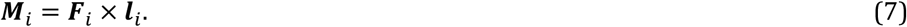

Now we calculate the friction torque *M*_*f*_. Let *ω* be the angular velocity of the body rotation about the COM. Then the torque of the friction force is

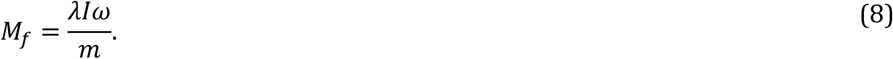

By Second Newton’s Law in angular form, after calculating the total torque we get the following differential equation describing the rotation of the body about the COM in the horizontal plane:

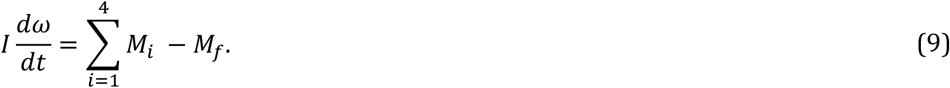

#### 2.3.2 Vertical forces

Here we calculate the vertical components of the ground reaction forces in each limb that we also refer to as weight-bearing or supporting forces. Let ***G***_*i*_ denote the supporting force in limb *i*. Then, 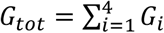 is the total supporting force. The weight distribution over the limbs depends on the number of supporting limbs, so below we discuss different possible situations depending on the number of limbs in stance.

##### Support by more than two limbs

We assume that the body frame always remains in the horizontal plane and, therefore, is not pitching or rolling. Therefore, the total torque created by the supporting forces should be balanced, i.e.

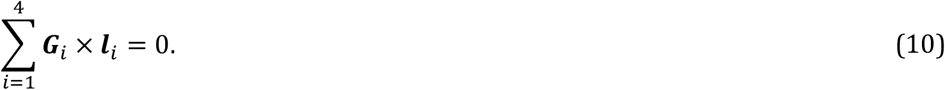

In addition, for the body to remain in the horizontal plane (not move in the vertical direction), the total supporting force should be equal in magnitude to the gravitational force:

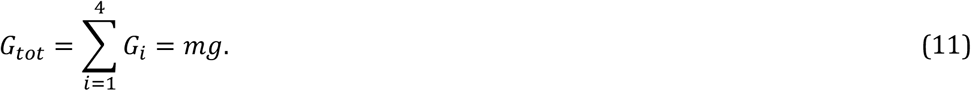

Finally, ground reaction forces are zero for all limbs in swing.

Note that Eq. (10) contains two equations as the limb displacements in the horizontal plane relative to the COM have two components. So, Eqs. (10) and (11) define a linear system of three equations for the supporting forces. This system has a unique solution in the case of three-leg support. In the case of four-leg support an additional constraint is necessary as different weight distributions that satisfy Eqs. (10) and (11) are possible. For four-leg support, we assume that the total load is distributed as evenly as possible. Particularly, we find a solution of Eqs. (10) and (11) that minimizes 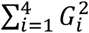 using the method of Lagrange multipliers.

##### Support by two limbs

In case of only two limbs being on the ground, the system of Eqs. (10) and (11) does not generally have solutions (unless the COM is precisely above the line of support, see Fig. 3). Here we take the case of the diagonal limbs in stance as an example. Consider the left hind (*i* = 3) and right fore *(i* = 2) limbs are supporting and the other two limbs (*i* = 1 and 4) are in swing (as shown in Fig. 3), that is, *G*_1_ = *G*_4_ = 0. The movement of this frame in the direction perpendicular to the supporting line will follow the inverted pendulum dynamics as illustrated in Fig. 3. Let *v*_*p*_ be the COM velocity component in the direction perpendicular to the supporting line. By taking the centripetal force into account, for the vertical component of the ground reaction force we have:

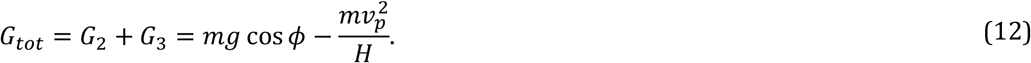

To find the load distribution over limbs 1 and 4, we assume that there is no pitch in the direction of the supporting line, so the torque created by *G*_2_ and *G*_3_ about the projection of the COM onto the line of support must be zero. Specifically,

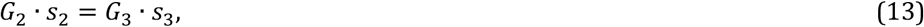

where *s*_2_ and *s*_3_ are the segments of the line of support between the projection of the COM on it and the positions of paws 2 and 3, respectively (see Fig. 3). After solving the linear system (12), (13) we find *G*_2_ and *G*_3_ for the case that the body is supported by limbs 2 and 3 only. Similarly, we find the vertical components of the ground reaction forces (also referred to as weight-bearing forces and limb load) for any other pair of supporting limbs during 2-leg support phases.

## 3 Results

Below, we introduce the locomotion control mechanisms, including those that define the stance-to-swing and swing-to-stance phase transitions. We then describe and explain the results of simulations based on different sets of assumptions. In our study, we focus on the locomotor regimes that result in the body moving in a straight-line. In the case of slow locomotion, the straight-line movement occurs when the gait is symmetric, i.e., when the left and right steps of contralateral limbs occur in exact anti-phase. Therefore, we investigate numerically and, where possible, analytically, the existence and stability of the symmetrical regimes in different model configurations and compare their characteristics with available experimental data on freely walking mice.

### 3.1 Transitions between swing and stance

A limb has two states during movement, swing and stance. When the limb is in the stance phase, as we described previously by decomposing the ground reaction force, it supports the body vertically and pushes it horizontally to move forward. When the limb is in swing, both the horizontal and vertical components of force are equal to zero and the limb instantaneously moves in the air to its target position relative to the body.

To organize the transition from stance to swing in our model, we followed the previous model of Ekeberg and Pearson [25, 32] and the rules formulated in that model. Based on these rules, there are two conditions for stance-to-swing transition: limb unloading (transferring the load to other limbs) and limb (over)extension. In animals, this transition is controlled by two sensory signals, the force-dependent feedback from ankle extensor muscles and the length-dependent feedback from hip and ankle flexor muscles. These two conditions have been implemented in our model (see Fig. 1B). First, if the vertical component of the ground reaction force of a limb (load) is not greater than 0, which means it does not support the body, the paw detaches from the ground, and that limb switches from stance to swing. Second, we assume that the displacement of the paw from its initial position in the coordinate system associated with the body is limited by the maximum limb displacement *D*, so that the limb in stance must transition to swing once its displacement reaches *D*. Therefore, the limb *i* is lifted if either of the following conditions is satisfied:

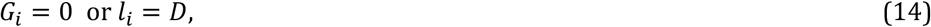

where *G*_*i*_ is the *i-*th limb load, *l*_i_ is the distance from the *i-*th limbs paw to its initial position relative to the body, and *D* is the maximum limb displacement (see Table 1 for parameter values).

**Table 1.** Model parameters.

| Name | Notation | Value |
| --- | --- | --- |
| Length of the animal | $L$ | 10 cm |
| COM height | $H$ | 3 cm |
| Half width | $h$ | 1 cm |
| Maximal limb displacement | $D$ | 5 cm |
| Mass | $m$ | 25 g |
| Friction coefficient | $\lambda$ | 100 g/s |
| Gravitational acceleration | $g$ | 1000 cm/s<sup>2</sup> |

For the transitions from swing to stance, for which the mechanisms are still elusive, we explored two possibilities: the direct control of swing duration (i.e., constant swing duration imposed by the central neural controller, see Model 1 described in section 3.2), and the control of swing termination through balance feedback signaling on postural instability of the body (Model 2 and Model 3, described in sections 3.3 and 3.4).

### 3.2 Model 1: Feedback independent swing duration

In Model 1, we assume that the termination of swing (and hence the swing-to-stance transition, or limb landing) is directly controlled by its (homonymous) rhythm generator within the spinal cord, which makes the swing duration constant (dependent only on the properties of the rhythm generator and independent of external inputs, such as feedback from the limbs). In other words, each limb moving down touches the ground at a fixed time interval *t*_*sw*_ after the start of the swing phase, which becomes a parameter of the model. In addition, Model 1 does not have any central interactions between the rhythm generators (see Fig. 1).

#### 3.2.1 Model 1 operation

Figure 4A shows the stable behaviors of Model 1 obtained by varying two parameters, the propulsion force *F*_0_, generated by each limb during stance, and the fixed swing duration *t*_*sw*_. Each point in the plot corresponds to one step cycle with the x-coordinate representing the swing duration (*t*_*sw*_), the y-coordinate representing the average COM velocity over the step cycle (i.e., COM displacement divided by the step cycle period), and the color indicating the duty factor (i.e., the stance duration divided by the step cycle period). Previously, by fitting experimental data on overground mouse locomotion, Herbin et al. [29] found empirical relationships between swing duration (*t*_*sw*_, s) and stride frequency (*f*, Hz): *t*_*sw*_ = 0.121 − 0.006 *f*, and between stride frequency (*f*, Hz) and velocity (*v*_*c*_, cm/s): *f* = 1.004 + 0.4 *v*_*c*_/ ln *v*_*c*_). We combined these approximations to express experimental swing duration as a function of locomotor velocity *t*_*sw*_= 0.115 − 0.0024 *v*_*c*_/ ln *v*_*c*_, which is shown in Figs. 4, 8, and 10 by a thick green line.

**Figure 4:**
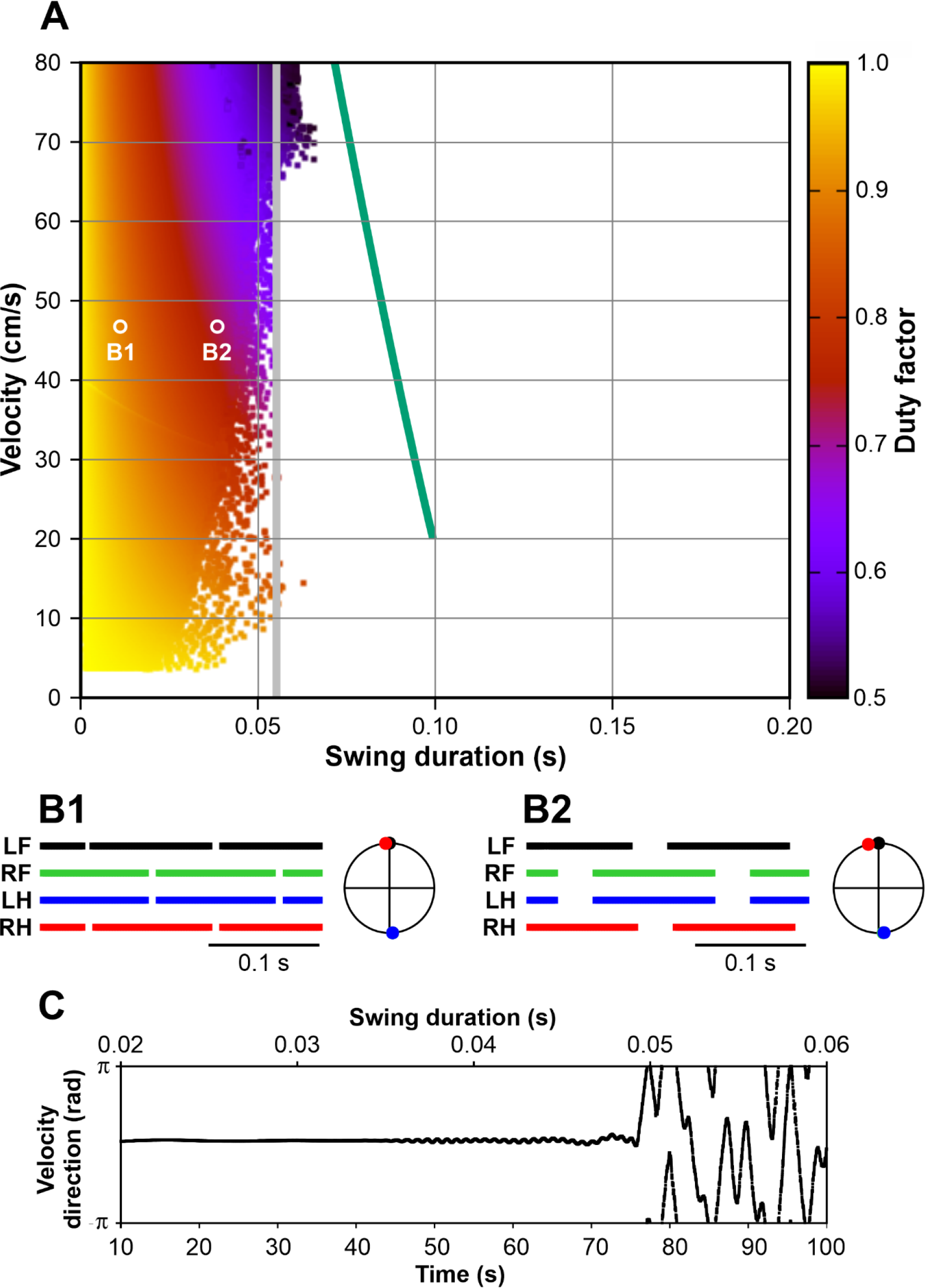
Symmetric gaits observed in Model 1. **A**. Heatmap representation of locomotion. This heat map was calculated based on 200 simulations for the force value *F*_0_ varying between 0 and 5,000 g·cm/s^2^ with a step of 250 g·cm/s^2^. For each force value, the model was integrated for 100 s with the swing duration parameter linearly changing from 0.001 to 0.1 s. The initial conditions were chosen so that all limbs were in stance, the diagonal limbs had the same phase, and left and right limbs were in anti-phase. Each point of the heat map represents one step cycle with the swing duration along the horizontal axis and the COM velocity averaged over the step cycle on the vertical axis. The color reflects the duty factor according to the color key on the right. The simulation was terminated if the relative cycle-to-cycle change in the duty factor (DF) exceeded a preset threshold (specifically, if | ln(*DF*_*current*_/*DF*_*previous*_) | > 0.2) which was used as an indicator that the tracked regime was destroyed or lost stability. We have also confirmed that once the regime loses stability according to this criterion, the subsequent model simulation eventually results in a fall, so no other stable gaits were observed. The vertical grey line shows the inverted pendulum time constant of 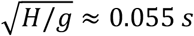 (see section 3.2.4). **B1 and B2**. Stepping diagrams (stance phases of all limbs) and phase relationships between the limbs at small (0.01 s) and relatively large (0.035 s) swing durations. In both regimes, the diagonal (LF-RH, RF-LH) limbs are synchronized (zero phase difference) and left and right limbs are in anti-phase, indicating a symmetric gait. **C**. Instantaneous movement direction in a model simulation at a locomotor velocity of approximately 47 cm/s with the swing duration slowly increasing throughout the simulation. Swing durations are indicated at the top. This corresponds to moving along a horizontal line through labels B1 and B2 in panel **A**. The direction of movement is constant until the swing duration reaches ∼0.035 s. After that slow oscillations emerge in the movement direction and the symmetric gait starts destabilizing. After the swing duration exceeds ∼0.05 s, the walking becomes highly unstable and eventually comes to a fall. See the supplemental video [model1.mp4](https://drive.google.com/file/d/1uZqTt_Iiji_V_qdKL2_o-wt80o9HdMpW/view?usp=drive_link).

Several major observations can be formulated from our simulations. First, stable locomotion in this model is represented by a symmetric gait in which diagonal limbs are fully synchronized (see details below in section 3.2.2) and left and right limbs are alternating (move in anti-phase, Fig. 4B1, B2). Second, since the diagonal limbs swing together, the body is supported either by two legs (during the two swing phases of the synchronous diagonal limbs) or by four legs (at all other times). This gait is characterized by a high duty factor and very short swing duration, which is unusual for mouse locomotion [31, 33]. Third, our simulations could not produce stable symmetric gaits with a swing duration greater than ∼0.055 s (Fig. 4A). This contrasts with swing durations in behaving mice that can vary between 0.07 s and 0.1 s depending on the locomotor velocity [29, 34, 35].

Below we explain the synchronization properties and the swing duration limitations of Model 1.

#### 3.2.2 Diagonal synchronization

Remarkably, Model 1 generates a very specific limb coordination pattern, despite the absence of central interactions between rhythm generators controlling the four limbs. One of the characteristics of this pattern is the synchronization of the diagonal limbs. The biomechanical reasons for this are as follows.

At a relatively short swing duration, the body is supported by four limbs most of the time. In this case, lifting one of the limbs can lead the COM to be outside the triangle formed by the remaining three limbs on the ground, which leads to immediate unloading of the limb diagonal to the lifted one (see Fig. 5A). For example, lifting the right hindlimb in the first example shown in Fig. 5A would immediately unload the left forelimb (since the COM is outside of the triangle formed by the limbs remaining on the ground) and thus trigger the liftoff of the left forelimb. In the second example, raising the left forelimb also immediately causes lifting of its diagonal counterpart (the right hind limb) due to unloading. Thus, at least for a subset of initial conditions similar to the examples shown, the diagonal synchronization of the limbs occurs within one step.

**Figure 5.**
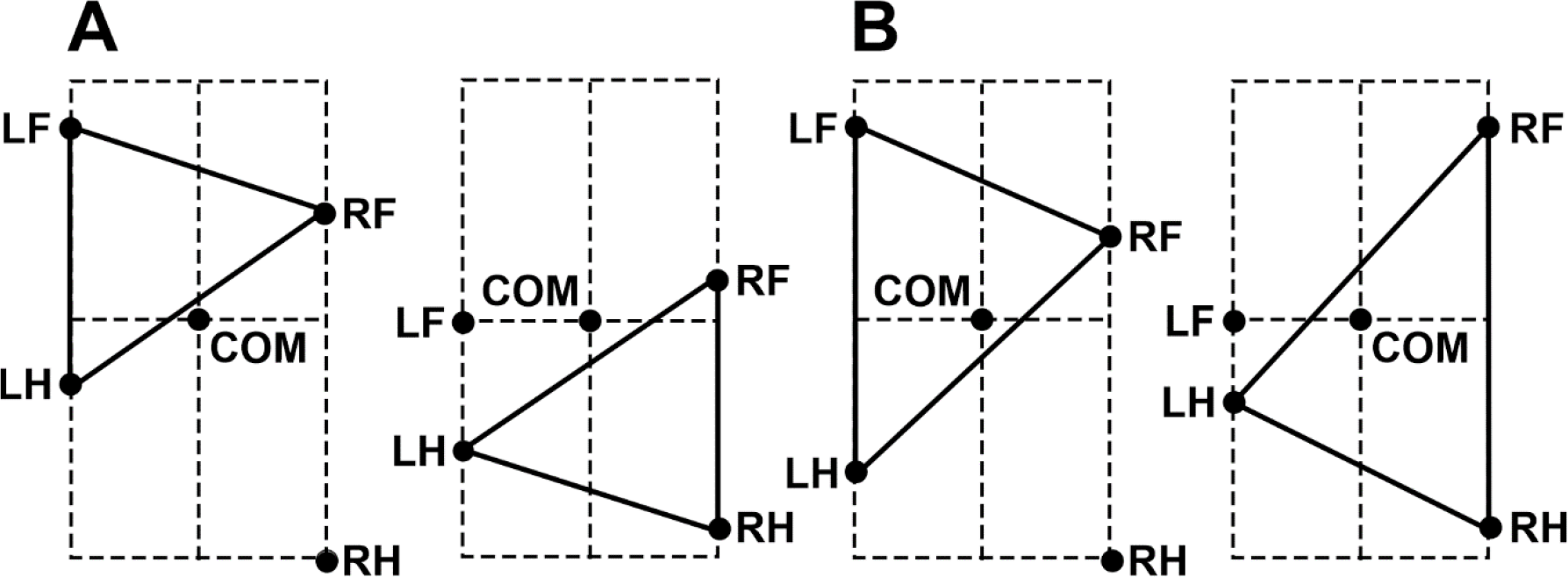
Diagonal synchronization in Model 1 due to limb unloading. **A**. On the left, the RH limb is fully extended and is about to lift off. After that, the COM appears outside of the triangle with vertices at the paws remaining on the ground, meaning that the LF is unloaded and, therefore, gets lifted immediately too. On the right, a similar situation is shown, where the liftoff of the LF limb causes the RH limb to immediately transition to the swing phase too. **B**. In these configurations, the COM remains within the supporting triangle after the fully extended limbs (RH on the left and LF on the right) transition to the swing phase, and no immediate diagonal synchronization occurs.

For other configurations (i.e., where the COM appears inside of the triangle formed by the limbs remaining on the ground, see Fig. 5B), another synchronization mechanism kicks in. The example shown in Fig. 6 shows the situation when all four limbs are supporting the body, but the first and the fourth limbs have reached their maximum extension and are about to be lifted. If we perturb the system so that it lifts the first limb a little bit before it reaches its maximum extension, the number of limbs supporting the body on the left and right sides changes. In our example, two limbs push forward on the right side while only one limb pushes on the left (see arrows in Fig. 6). This creates an uncompensated yaw torque, and the body starts rotating counterclockwise around the COM. As a result, the fourth limb extends faster than when the body does not rotate. Therefore, the fourth limb reaches its maximum length faster and is lifted earlier, thus decreasing the phase difference between the diagonal limbs in every step and gradually restoring synchronous movements of the diagonal limbs.

**Figure 6.**
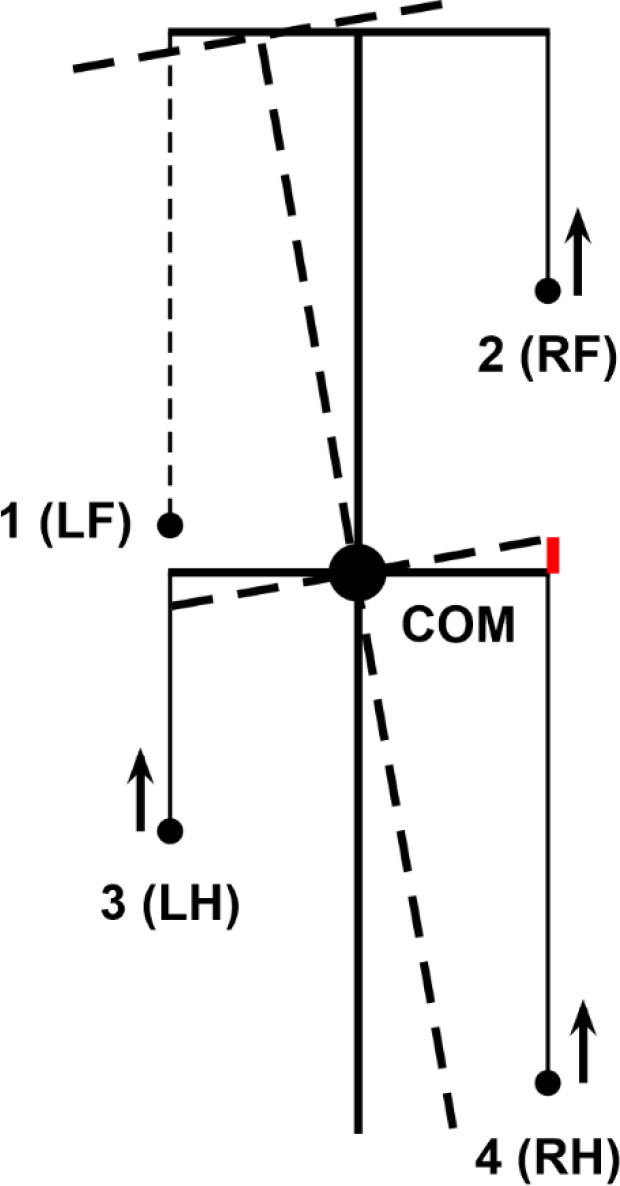
Synchronization of diagonal limbs by body rotation. Lifting the LF limb before the RH limb creates an uncompensated yaw torque that rotates the body counterclockwise (since there are two limbs pushing forward on the right side of the body, while there is only one on the left, see arrows). This rotation creates a faster extension of the RH limb compared to the unperturbed movement during which the body is oriented straight forward. The difference between the RH limb extension in the perturbed and unperturbed cases is indicated in red. Faster leg extension leads to an earlier transition of the RH limb into the swing phase, thus reducing the phase difference between the LF and RH limbs. Asymptotically, the synchronization between the LF and RH limbs is restored.

#### 3.2.3 Gait symmetry

As already noted, the gait in Model 1 is characterized by an interlimb coordination pattern, in which the two-limb-supporting periods alternate with four-limb-supporting periods and the swing phase of each limb occurs at the middle of the stance phase of the contralateral limb. Below we explain the mechanism that stabilizes this left-right alternation of the left and right steps occurring in exact anti-phase, which also results from the body rotating due to an uncompensated yaw torque.

Figure 7 shows the scenario when the body is first supported by all four limbs [see the first (leftmost) black segment of the trajectory], then LF and RH limbs are simultaneously lifted while RF and LH remain on the ground (see the dashed line connecting LH and RF limbs) supporting the body during the red segment of the COM trajectory. At the end of this segment, LF and RH limbs make a touchdown (as shown by the dashed line connecting LF and RH in Fig. 7) initiating the second epoch of the 4-leg support (see the black trajectory segment in the middle) which lasts until LH and RF are lifted off. This starts the second epoch of 2-leg support shown by the solid red segment of the trajectory crossing the LF-RH diagonal in Fig. 7.

**Figure 7.**
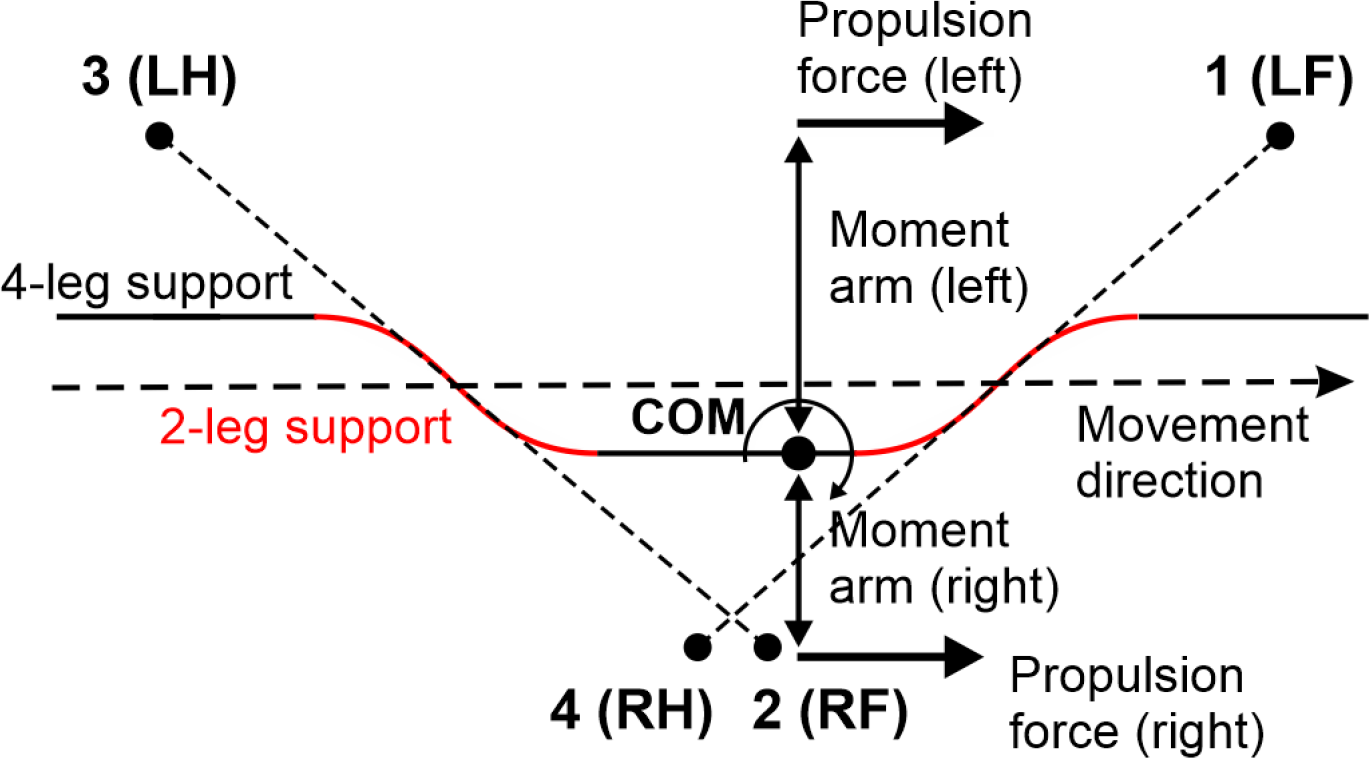
Schematic COM trajectory produced by Model 1. The gait is alternating 4- and 2-leg support phases. Segments of the trajectory during 2- and 4-leg support are shown in red and in black, respectively. Dotted lines show the lines of support during 2-leg support phases during which the trajectory bends (red segments). As a result, during movement, the COM displaces from the (dashed) midline to the left or right, creating unequal moment arms for the propulsion forces applied on different sides. Unequal moment arms lead to non-zero net yaw torque which rotates the body about the COM during 4-leg support phases. See text for more details.

**Figure 8.**
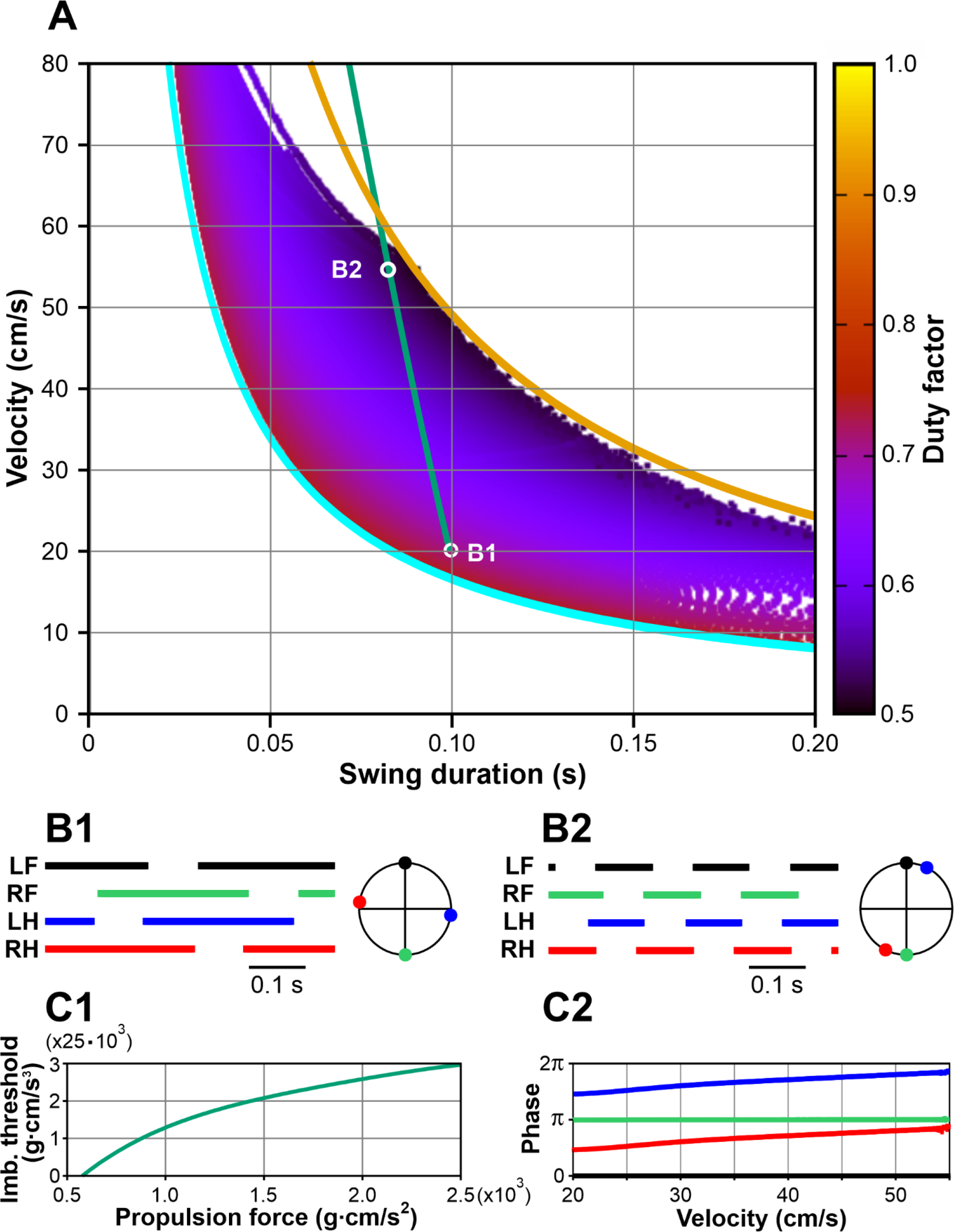
Symmetric gaits observed in Model 2. **A**. Heatmap representation of locomotion. This representation was constructed similar to the one shown in Fig. 4A, but instead of slowly changing the swing duration at fixed values of the propulsion force, we progressively increased the threshold for the imbalance signal *k*_*im*_ from 0 to 250,000 g·cm/s^3^, which resulted in a corresponding increase in swing duration. Each point of the heatmap represents one step cycle, x-coordinates represent swing durations, y-coordinates represent average velocities, and the color reflects the duty factors according to the color key on the right. At zero threshold *k*_*im*_ = 0, Model 2 exhibits a lateral walking gait with swing durations equal to ¼ of the step cycle, corresponding to the maximal possible duty factor of 0.75. This corresponds to the left/lower boundary of the region when *v*_*c*_ *t*_*sw*_ = *D*/3 shown by the thick cyan curve (see text for more details). Due to the mentioned modeling constraints, the minimal possible value of the duty factor is 0.5, and the corresponding right/upper boundary of the region is *v*_*c*_ *t*_*sw*_ = *D* (shown by the thick orange curve). At locomotor velocities below 60 cm/s, with increasing *k*_*im*_, the gait exhibited by Model 2 continuously transforms from a lateral walk to a pace-like gait (with homolateral limbs swinging more and more synchronously) as the duty factor approaches 0.5. **B1** and **B2**. Stepping diagrams (stance phases of all limbs) and phase relationships between the limbs corresponding to regimes B1 and B2 in panel A. In both regimes, left and right limbs are in anti-phase indicating symmetry of the gait. At the lower velocity (**B1**), limb phases are evenly distributed over the step cycle consistent with a lateral walking gait. At the higher velocity (**B2**) the phases of homolateral limbs become very close indicating the transition to a pacing gait. **C1**. Model 2 reproduces the dependence of the swing duration on velocity as characterized by Herbin et al. [29] (the green line in panel **A**) as long as the imbalance threshold (*k*_*im*_) and the propulsion force (*F*_0_) parameters follow the relationship shown. **C2**. Limb phases relative to LF as the propulsion force and the imbalance threshold are varied as shown in C1, so that the swing duration and velocity change along the green line in panel **A** between the points labeled B1 and B2. Color coding as in B1 and B2. At the lowest speed, the phases correspond to a lateral walk (see **B1**). As speed increases, phase differences between homolateral limbs decrease, showing a gradual transition to pace (see **B2**). See the supplemental video [model2.mp4](https://drive.google.com/file/d/1Bpm9vnh6y9E5zGzOHk_7G9HcRU86Jke2/view?usp=drive_link).

During the two limb support phases, the COM motion is affected by gravity (see Eqs. (1)-(4)) which bends the COM trajectory as shown by the red segments in Fig. 7. Therefore, after every 2-leg support phase, the COM gets slightly shifted from the midline towards the left or the right side depending on the supporting diagonal. In Fig. 7 one can see that when the body is supported by the RF and LH limbs, the COM shifts to the right, and when the body is supported by the LF and RH limbs, the COM shifts to the left. During the subsequent 4-leg support phases the COM remains shifted towards one of the sides, which creates different moment arms relative to the COM for the propulsion forces created by the left and right limbs. For example, during the 4-leg support phase in the middle, the COM is shifted to the right side, and the moment arms of the left limbs’ propulsion forces are greater than the moment arms of the right limbs’ propulsion forces. Therefore, the propulsion forces applied on the left side create greater yaw torque than the ones applied on the right side which leads to clockwise rotation of the body about the COM as it moves along this part of the trajectory (opposite to the one shown in Fig. 6).

Let’s now consider a perturbation of the exact anti-phase alternations of the left and right swings, in which the liftoff of the LH-RF diagonal in Fig. 7 is slightly delayed. As a result, the 4-limb support phase in the middle is prolonged. Due to the longer duration of this phase, the body would rotate more in the clockwise direction which would delay the liftoff of the RH limb after the perturbation based on the mechanism described in the previous section (if the body frame rotates clockwise in Fig. 6, the RH extension slows down instead of speeding up and its liftoff gets delayed). This delay partially compensates for the perturbation effect and asymptotically realigns the swings of the RF-LH and LF-RH diagonals in exact anti-phase and thus restores gait symmetry.

#### 3.2.4 Range of possible swing durations

It should be noted that the synchronization mechanism described above is rather weak and requires multiple steps for the system to converge to the symmetric gait. Consequently, any destabilizing factor with a relatively short timescale can destroy this regime. The most obvious instability in this model is associated with the inverted pendulum dynamics that has a time constant of 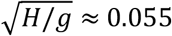, where *H* is the COM height and *g* is the gravitational acceleration (see Eq. (1)). In gaits exhibited by this model, the body movement during swing is described by the inverted pendulum dynamics. Therefore, it is reasonable to expect that if the swing duration is comparable or exceeds 0.055 s, the symmetric gait can destabilize. This is supported by our simulations. As seen in Fig. 4A, the range of possible swing durations for stable locomotion does not extend beyond 0.055 s (see the vertical gray line). This limitation of swing duration clearly contradicts experimental data on overground and treadmill walking in mice which always exceed 0.07s and can be as long as 0.1 – 0.125 s [29, 34, 35]. The development of the symmetric gait instability in this model as the swing duration increases is illustrated in Fig, 4C where we show the changes in the velocity direction when the swing duration increases throughout the simulation. Movement direction is relatively constant at swing durations below approximately 0.035 s. A further increase of swing duration results in slow oscillations accompanied by body rotation (see the supplemental video [model1.mp4](https://drive.google.com/file/d/1uZqTt_Iiji_V_qdKL2_o-wt80o9HdMpW/view?usp=drive_link)). When the swing duration reaches about 0.05 s, these oscillations destabilize, and the movement direction starts changing uncontrollably. This results in a misalignment of the body with the movement direction and eventually leads to circling movements, tripping, and falling (see the video [model1.mp4](https://drive.google.com/file/d/1uZqTt_Iiji_V_qdKL2_o-wt80o9HdMpW/view?usp=drive_link)). Based on this, one can speculate that the gait instability originates from complex interactions between the COM displacement in the frontal plane, body rotation due to the emerging uncompensated torque, and the mechanical feedback (i.e., limb loading and extension). Even though there is a mechanism that maintains the symmetry between left and right legs (i.e., alternation of left and right steps in exact anti-phase) at relatively short swing durations (see section. 3.2.3), this mechanism fails once the swing duration reaches values comparable to the inverted pendulum time constant, which is significantly smaller than typical swing durations observed in mouse locomotion. It is important to note that it would be impossible to study gait instabilities resulting from interactions between movements of the body in the frontal plane, body orientation, and sensory feedback in a model describing one-dimensional movements only. To the best of our knowledge, the formalism we are presenting in this study is the first mathematically tractable model suitable to study locomotion on a plane.

#### 3.2.5 Possible involvement of central interactions

It is important to highlight that Model 1 does not have neural mechanisms within the central controller that could contribute to the phase coordination between RGs controlling different limbs. Different types of such central interactions between the RGs, particularly those involved the left-right inhibition, diagonal excitation, or homolateral inhibition, were included in our previous models [5-7, 36-40] to meet and/or reproduce multiple experimental data [2, 3, 39-43]. Therefore, we checked the idea that incorporating such central interactions would allow Model 1 to locomote with longer (more realistic) swing durations while supporting the left-right symmetry necessary for maintaining the direction of movement. However, incorporating the above-mentioned central interactions in Model 1 had no effect on the generation of symmetric gaits and their stability. Therefore, these central interactions were unable to extend the range of possible swing durations. An explanation of this failure is provided below.

The functional role of left-right inhibition between flexor/swing half-centers is to prevent them from being active at the same time. However, in the gait we characterized in Model 1, the left and right swings always occur in antiphase and, therefore, never overlap. Thus, additional left-right inhibition cannot have any significant effect on the produced gait, as the contralateral inhibitory signals arrive during the silent phase of the flexor/swing half-centers and cannot alter their activity. Similarly, homolateral inhibition should prevent the flexor/swing half-centers on the same side of the body from being simultaneously activated. However, due to the diagonal synchronization of limbs in Model 1, fore- and hind limbs on the same side of the body alternate in antiphase in the same way as left and right limbs. Therefore, the addition of homolateral inhibition has no consequences for the existence or stability of the observed gait. Finally, the role of diagonal excitation is to synchronize the movements of the diagonal limbs. As extensively analyzed in section 3.2.2, Model 1 exhibits robust diagonal synchronization solely based on interlimb interactions mediated by biomechanics. This is why the inclusion of direct interactions between diagonal flexor half-centers has no additional effect.

### 3.3 Swing duration defined by balance control. Models 2 and 3

#### 3.3.1 cLoss of balance

As described above, the exclusively feedforward control of swing duration (independent of external factors, of central interlimb coordination pathways, and of the system’s interactions with the environment) implemented in Model 1, was unable to support stable straight-line locomotion with swing durations that reach experimentally observed values. This suggests that, although the intrinsic dynamics of rhythm generators may contribute to swing duration control, they might not alone define the swing duration and gait expression seen in mouse locomotion. Therefore, we have considered two possible external signals that can trigger the swing-to-stance transitions: 1) feedback signals representing a loss of balance and 2) informing the central controller of a critical loss of balance.

Loss of balance in our model can occur during 2-leg support phases as the COM is dragged from the supporting line by gravity. As this happens, the total weight-bearing force (a sum of the vertical components of all ground reaction forces) decreases. However, the value of the total load itself cannot be used as a reliable signal indicating loss of balance, as the ground reaction forces are reduced by the centrifugal force which increases with COM velocity (see Eq. (12)). Therefore, we define loss of balance in our model as the event when the total supporting force decreases faster than at a certain rate. Specifically, we introduce an *imbalance threshold k*_*im*_ ≥ 0 so that the model mouse is losing balance when

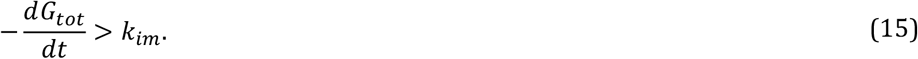

We refer to the left-hand side of this inequality as an *imbalance signal*. Notice that this is a global signal received by all four rhythm generators, which is different from the individual swing duration control that occurs independently in each limb. Therefore, when more than one limb is in swing, it must be decided which of them should transition into stance. We simply assume that when an imbalance occurs based on (15), the limb swinging for the longest time immediately switches to stance.

#### 3.3.2 Model 2. Swing-to-stance transition based on balance control

In contrast to Model 1, the duration of swing in each limb in Model 2 was not fixed (i.e., was not defined centrally within the corresponding RG). Instead, the timing of the swing-to-stance transition was defined by the common imbalance signal reaching the threshold *k*_*im*_ (see Eq. (15)). To characterize possible behaviors of Model 2, we used the same representation as in Model 1. We swept parameters *k*_*im*_ and *F*_0_ and calculated the corresponding COM velocity and swing duration. The results are shown in Fig. 8A where we also represent experimental data from mice during overground locomotion (green line, as in Fig. 4A). Our simulations show that Model 2 can generate stable symmetric gaits with considerably longer swing durations when compared to Model 1 and exhibits physiologically realistic swing durations for velocities between 20 and 55 cm/s.

### 3.3 Gait characteristics in Model 2

In Model 2, each swing phase is terminated when a postural imbalance occurs, which can only happen during 2-leg support phases. Once one of the two swinging legs touches down, the other will remain in swing until the balance is lost again, which can only happen after one of the three supporting legs lifts off. This implies that in Model 2 all four limbs will never be on the ground at the same time, i.e., at least one limb will be in swing at any given time. As shown in Fig. 8A, the swing duration in Model 2 has a lower boundary defining a velocity-dependent minimum swing duration that corresponds to a duty factor of 0.75. This boundary represents the extreme case, when there is exactly one limb swinging while three other limbs remain on the ground (the step cycle is divided into exactly four swing phases, meaning that the stance phase duration for each leg is equal to three swing durations). During stance, the displacement of the paw relative to its initial position in the body coordinate system is equal to the velocity multiplied by the stance duration. The stance is terminated when this displacement reaches *D*. Taken together, one can write the following equation for the minimal swing duration (corresponding to the cyan curve in Fig. 8A):

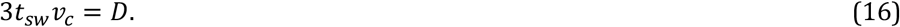

The swing duration also has an upper boundary defining a velocity-dependent maximum swing duration that corresponds to the duty factor of 0.5, when there are 2 limbs on the ground at all times. This boundary (orange curve in Fig. 8A) reflects a general limitation of our modeling framework: since there may not be more than two limbs simultaneously in swing, in the extreme case the step cycle period would be exactly equal to two swing durations. In such a regime, the stance duration is equal to the swing duration, so the equation for the upper boundary is

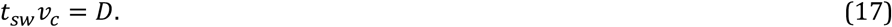

#### 3.3.4 Symmetric gait robustness

Our simulations show that as we increase the imbalance parameter *k*_*im*_ from zero up (see Eq. (15)), Model 2 exhibits a stable symmetric gait almost everywhere between the two boundaries described by Eqs. (16) and (17) (see cyan and orange curves in Fig. 8A) except for velocity values above 55 cm/s. This velocity roughly corresponds to the Froude number 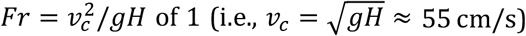 which defines the maximal possible walking speed [44]. Faster running includes suspension phases that our modeling framework does not describe. The stability of the symmetric gait below 55 cm/s is achieved due to an extreme robustness of the exact anti-phase left-right synchronization of the limbs. As illustrated in Fig. 9, touchdowns of the swinging limbs occur at very specific configurations of the supporting legs relative to the COM which leads to virtually instantaneous adjustments in response to perturbations, so that the symmetry of the gait gets restored within a single step cycle. This is in contrast with Model 1 where left-right anti-phase synchronization relies on a much weaker mechanism which eventually fails to counteract the instability in movement direction that occurs after the swing duration becomes comparable to the inverted pendulum time constant (see Sec. 3.2.4, Fig. 4C and the corresponding supplemental video [model1.mp4](https://drive.google.com/file/d/1uZqTt_Iiji_V_qdKL2_o-wt80o9HdMpW/view?usp=drive_link)).

**Figure 9.**
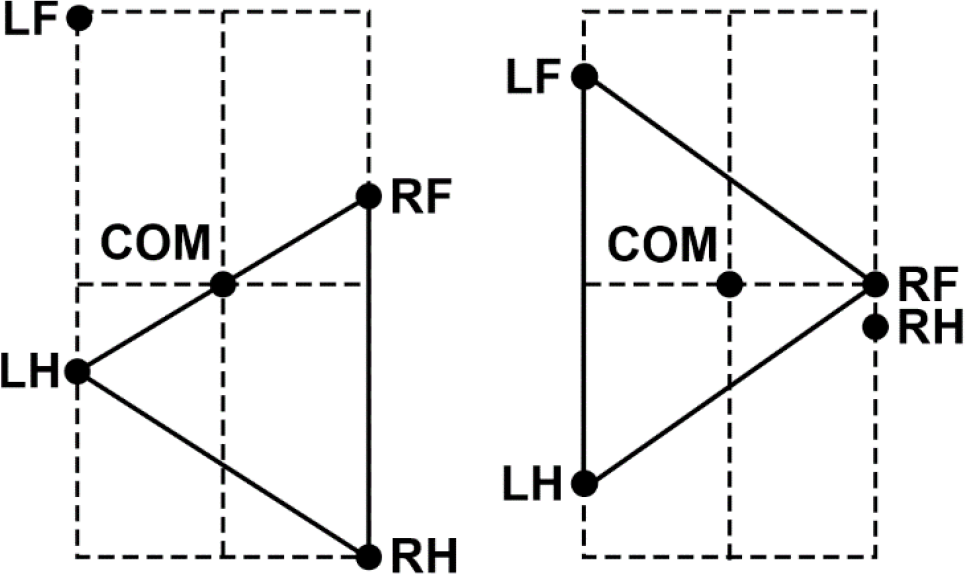
Swing to stance transitions in Model 2. **Left:** when RH is lifted due to unloading/full extension, the COM crosses the LH-RF supporting line and starts falling, causing LF to land. **Right:** when RF lifts off due to full extension, the COM starts falling to the right causing RH to land. This creates the limb movement sequence LF-RH-RF-LH, which corresponds to a lateral walking gait.

**Figure 10:**
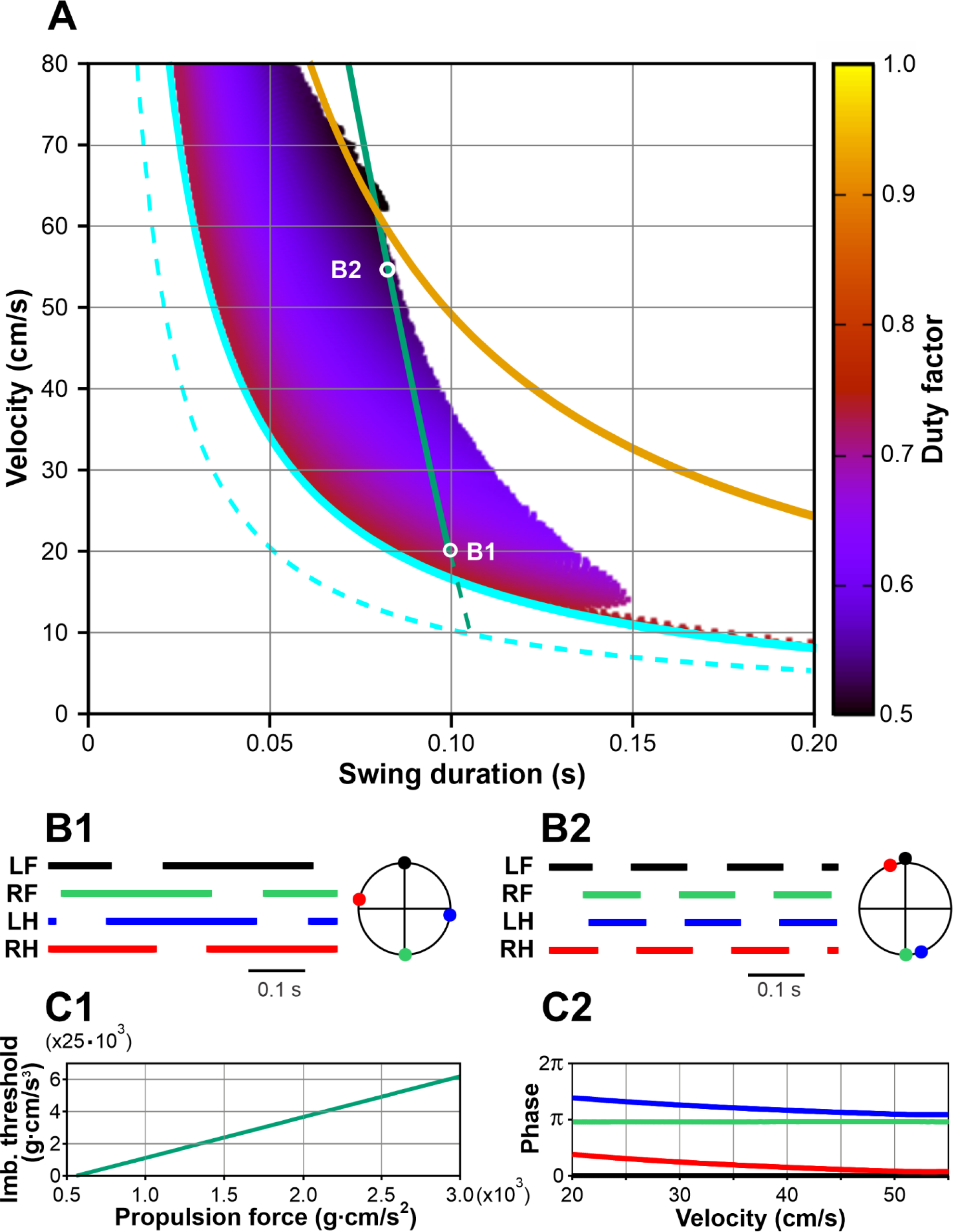
Symmetric gaits observed in Model 3. **A**. Heatmap representation of locomotion. Each point represents a step cycle with the swing duration on the x-axis, the average COM velocity on the y-axis, and the duty factor in color according to the color key on the right. As for Model 2, possible gaits lie between the curves corresponding to the duty factors of 0.75 (the lower boundary, solid cyan curve) and 0.5 (the upper boundary, yellow curve). For velocities below 60 cm/s, the swing duration becomes bounded by increasingly stronger lateral COM oscillations. The green line, representing experimental data [29] in this figure as well as in Figs. 4 and 8, was extrapolated to lower velocity values (see its dashed segment) to include locomotor behaviors at lower speeds. The dashed cyan curve corresponds to a reduced value of the maximal limb displacement (*D* = 3 cm). See text for details. **B1 and B2**. Stepping diagrams and phase relationships between the limbs near the lower boundary (B1) and close to the upper boundary (B2). **C1**. The relationship between the imbalance threshold (*k*_*im*_) and the propulsion force (*F*_0_) in Model 3 that provides the experimentally observed dependence of the swing duration on velocity as in Herbin et al. [29] (along the green line in panel A). **C2**. Phases of the limbs relative to LF as the propulsion force and the imbalance threshold are varied as shown in C1. At the lowest speed the phases correspond to lateral walk (see **B1**). As the speed increases, phase differences between diagonal limbs reduce, showing a gradual transition to trot (see **B2**). See the supplemental video [model3.mp4](https://drive.google.com/file/d/13hx0M9_-nZ4EMQUPwU0UpOoUyGuHAeD/view?usp=drive_link).

#### 3.3.5. Transition from lateral walk to pace

The value of *k*_*im*_ = 0 corresponds to the case with minimal possible swing duration (see Eq. (20)). The COM moves forward, generally unloading the hind limbs and increasingly loading the forelimbs. When one of the forelimbs is in swing (e.g., LF, see Fig. 9 left), the first limb to unload is the diagonal hind limb (RH). This happens when the COM is exactly above the line connecting LH and RF. After RH lifts off, the COM crosses that line which triggers the imbalance feedback and causes LF to immediately land thus creating a new supporting triangle. Eventually, one of the forelimbs gets fully extended (RF in Fig. 9 right) and lifts off. At this time, the hindlimb on the same side is in swing (RH in Fig. 9 right). After that, the body is supported by the left (LH and LF) limbs only, which makes it fall to the right causing the RH limb to make an immediate touchdown. The step cycle continues similarly for the two remaining limbs thus creating a limb movement sequence of LF-RH-RF-LHF representing a lateral-sequence walk (Fig. 8B1).

At *k*_*im*_ > 0, the transition from swing to stance does not occur right after the body starts losing support, but only after the rate of change of the total load becomes large enough (see Eq. (15)). Therefore, swing phases of both diagonal and homolateral limbs start overlapping, and the two emerging overlapping intervals increase with the increase of *k*_*im*_. However, the overlap of the diagonal swing phases increases substantially slower than the overlap of the homolateral swing phases. This phenomenon has non-trivial mechanics. When two diagonal limbs are in swing, the body is supported by two other diagonal limbs. During such a phase the COM movement is affected by gravity pushing the COM in the direction perpendicular to the line of support. In the example shown in Fig. 9 Left, once RH lifts off, the COM trajectory starts bending to the left, which creates a velocity component towards the left-hand side. Importantly, the longer the duration of this phase is, the more the COM shifts to the left, and the greater the component of the velocity perpendicular to the direction of movement becomes. After the LF limb lands, the body gains 3-leg support, during which the COM continues shifting to the left, so that at the time when the RF limb lifts off, the COM is close to the LF-LH line of support and has significant initial velocity towards the left-hand side. For the imbalance feedback to kick in, the total ground reaction force must start falling. For this to happen, the COM movement to the left must be stopped by gravity first. The magnitude of this “inverted pendulum” force is proportional to the distance from the COM to the line of support, which becomes smaller as the COM shifts to the left-hand side. As both the shift and the velocity of the COM in the frontal plane increase with the imbalance threshold, the time spent before the COM reverses from moving to the left to moving to the right quickly grows, which leads to a progressively longer overlap of homolateral swings. In fact, the duration of the homolateral support phase increases much faster than the duration of the diagonal support with the increasing imbalance threshold. The resulting gait, as the swing duration approaches its upper boundary, becomes close to pace [45] (Fig. 8B2 and the supplemental video).

#### 3.3.6 Comparison with experimental data

In terms of the relationship between the locomotor velocity and the swing duration, Model 2 is compatible with mouse locomotion where the green line in Fig. 8A (representing the experimentally observed dependence [29]) lies inside of the region of stable symmetric gaits. This relationship is reproduced by the model given that the propulsion force *F*_0_ and the imbalance threshold *k*_*im*_ follow the curve shown in Fig. 8C1. However, the gait demonstrated by Model 2 features a long 2-leg support by homolateral limbs which is characteristic for pace [45] (Fig. 8B2, C2), whereas experimentally mice mostly exhibit trot and not pace [30, 31]. In trot, diagonal limbs move synchronously so that homolateral limbs alternate in the same way as left and right limbs do. To address this inconsistency, we considered Model 3, which represents a modified version of Model 2 that includes additional interlimb coordination mechanisms favoring diagonal synchronization.

### 3.4. Balance control and central mechanisms for interlimb coordination

As noted above, Model 2, which uses imbalance feedback to control swing durations but does not have central interactions between the RGs controlling individual limbs, exhibits gaits with homolateral limb synchronization that resemble pace rather than trot at intermediate speeds. This suggests that coordination between the limbs is not exclusively dependent on the movement mechanics but likely involves some additional mechanisms. Central interactions between the spinal RG circuits were previously characterized both experimentally and computationally [5-7, 30, 37, 39-41, 46]. One of such interactions, the diagonal excitation between the flexor half-centers of the limb RGs is particularly interesting [7, 40] because it facilitates diagonal limb synchronization and thus, prevents homolateral limbs from swinging at the same time. Therefore, these central interactions can potentially prevent the expression of pace in the model and make trot the dominant gait, as is the case in rodent locomotion [28, 29]. These central neural interactions have been incorporated into Model 2 to generate Model 3.

Excitatory interactions between the RGs controlling diagonal limbs, particularly between their flexor half-centers, can facilitate the synchronous transition of these RGs from stance to swing. These diagonal excitatory connections from lumbar circuits (controlling hindlimbs) to cervical circuits (controlling forelimbs) are thought to be mediated by propriospinal commissural V3 neurons [40]. Therefore, we constructed Model 3 with the assumption that the swing-to-stance transitions in the forelimb RGs are modulated by diagonal excitation (see Fig. 1B) such that the imbalance threshold for the swing-to-stance transition for each forelimb is increased when the corresponding diagonal hindlimb is in swing. Specifically, in this model, the imbalance threshold *k*_*im*_ for a limb to switch from swing to stance is equal to 0 when the corresponding diagonal limb is in stance, or a preset positive value when the diagonal limb is in swing. We use this value as a control parameter and investigate possible gaits of Model 3 as we vary it analogously to what we did for Models 1 and 2. The results are presented in Fig. 10. Similar to Model 2, possible gaits here are bounded by the curves corresponding to duty factors of 0.75 and 0.5 (Fig. 10A). Note that Models 2 and 3 are identical at zero imbalance threshold, as is their behavior at the lower boundary, where both models exhibit a lateral walking gait with one limb swinging at a time (Fig. 10B1). With an increase in the imbalance threshold, the swing phases of the diagonal limbs start overlapping like in Model 2. However, unlike in Model 2, no large overlap of the homolateral swings occurs. As the imbalance threshold grows, the overlap of the diagonal swings progressively increases and the phase difference between the diagonal limbs reduces (Fig. 10C2). This creates a gait that converges to trot as the duty factor decreases (Fig. 10B2). The striking difference between the gaits observed in Models 2 and 3 is that as the imbalance threshold increases, Model 2 transitions to a pace while Model 3 transitions to a trot (Figs. 7B2, 9B2 and supplemental videos). This transition in Model 3 occurs because modulation of the imbalance threshold by excitatory inputs from diagonal limbs affects swing termination differently in fore and hind limbs. Indeed, using the left panel in Fig. 9 as an illustration, we can observe the following sequence of events. As the RH limb is lifted, it triggers an excitatory input to the LF limb that is currently in swing. This input increases the LF limb’s imbalance threshold, which in turn extends its swing duration. Consequently, this leads to an increased overlap in the LF and RH swing phases, effectively reducing the phase difference between these two limbs. In contrast, when the RF limb lifts off due to full extension (Fig. 9 Right), the RH limb’s imbalance threshold remains zero because the LF limb is on the ground and does not provide diagonal excitation to RH. Therefore, as soon as the RF limb lifts off, the COM begins to shift to the right due to gravity. This immediately triggers the transition of the RH limb into stance and, unlike in Model 2, no large overlap of the RF and RH swing phases occurs.

Our study primarily examined mouse locomotion at velocities exceeding 20 cm/s, as described by Herbin et al. [29] and Mendes et al. [35]. For these velocities, we established a constant maximum limb displacement (*D*) of 5 cm, resulting in a stride length that varied from 7.0 to 9.5 cm (see Fig. 11C). However, it is important to note that mice can also walk with slower velocities ranging from 5 to 20 cm/s [30, 31]. To achieve these slower velocities without changing gait, mice significantly decrease their stride length from approximately 7 cm to about 2.5 - 4 cm [30, 31]. To accommodate this data in our model, we reduced the parameter *D* to 3 cm. This adjustment shifted the lower boundary towards lower velocities (see dashed cyan curve in Fig. 10A) so that experimentally reported swing durations corresponding to these slower walking regimes are also reproducible by the model.

**Figure 11.**
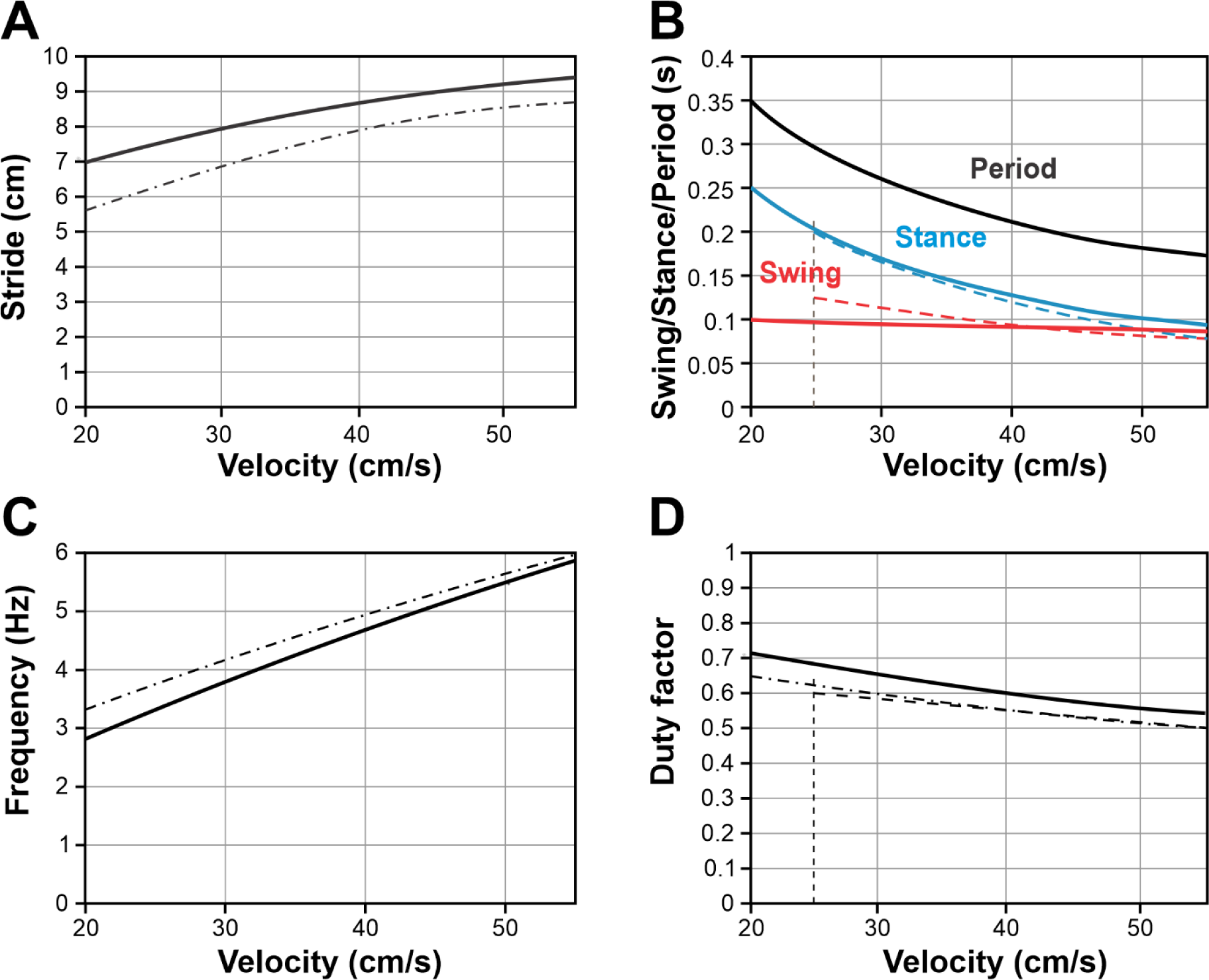
Model 3 performance in comparison with experimental data. **A-D**. The dependence of stride length (**A**), swing duration (**B**, red), stance duration (**B**, blue), step cycle period (**B**, black), stepping frequency (**C**), and duty factor (**D**) on velocity as generated by the model when the imbalance threshold and the propulsion force are varied along the line shown in Fig. 10C1. Despite the simplicity of our model, these relationships are in good qualitative correspondence with the experimental data [29, 35]. Solid lines show model results. Dashed lines in **B** and **D** are redrawn from Figures 3E and 3F of Mendes et al. [35]. Dash-dotted lines in **A, C**, and **D** are redrawn from Figures 1A and 3 of Herbin et al. [29].

### 3.5 Model 3 is consistent with mouse locomotion

As described above, Model 3, which includes balance control as a mechanism of swing termination and excitatory interactions between the diagonal RGs during their swing states, can reproduce the experimentally observed relationships between swing duration and locomotor velocity as well as the corresponding gaits exhibited by mice. To further evaluate the model, we examined whether other locomotor variables, such as stride length, stance duration, stepping period, stepping frequency, and duty factor, would align qualitatively with experimental observations.

To compare the model performance with the experimental data, we first characterized the relationship between the imbalance threshold (*k*_*im*_) and the propulsion force (*F*_0_) that ensures the dependence of the swing duration on the velocity during mouse overground locomotion published by Herbin et al. [29] (see the green line in Figs. 4A, 8A, 10A). This relationship appears to be linear as shown in Fig. 10C1.

We then varied the imbalance threshold *k*_*im*_ and the propulsion force *F*_0_ along this linear relationship and calculated the corresponding locomotor velocity, stride length, stance duration, step cycle period, stepping frequency, and duty factor. The graphs of all these characteristics and their changes with velocity are shown in Fig. 11 together with the analogous experimental dependencies drawn from [29] and [35] for a range of locomotor speeds from 20 to 55 cm/s. Taking into account the extreme simplifications assumed by the model, our model simulations and the experimental data are in good qualitative agreement.

## 4 Discussion

### 4.1 Rhythm generator as a “state machine”: state transitions, and sensory feedback

The concept of ‘*central pattern generators’* (CPGs), neural networks capable of generating rhythmic activity without rhythmic inputs, as the fundamental source of rhythmic motor behaviors such as locomotion, was formulated in early work by Graham Brown [47] and further developed and elaborated in many later studies [1, 4, 5, 8, 48-53]. The ability of neural circuits in the mammalian spinal cord to autonomously generate a locomotor-like rhythmic pattern of activity has been demonstrated both *in vivo* in decerebrated immobilized animals [51, 54-56] and *in vitro* in isolated spinal cords of rodents [2, 12-16, 57, 58]. This common locomotor-like pattern includes several rhythmic components associated with (and controlling) each limb and having specific phase relationships with each other. Particularly, the most typical pattern observed during fictive locomotion exhibits alternation between the flexor and extensor activity of the same limb and between the corresponding (flexor or extensor) activity of the left and right limbs [1, 2, 4, 8, 12-16, 54, 56, 57]. In the present study, we consider the CPG (or central controller) as the entire locomotor network controlling all limbs and assume that this CPG consists of four interacting ‘rhythm generators’ (RGs), each controlling the movement of one limb. While functional types of synergistic groups of muscles in each limb operate at different phases across the locomotor cycle, the output of each RG more broadly represents alternating flexor and extensor phases. Moreover, each RG could be considered as a neural structure with a dual function: 1) as a ‘*state machine*’ [59-61], defining the state and operation of the controlled system (limb) in each phase, and 2) as an ‘*oscillator*’, defining phase transitions between the states/phases (with or without external signals controlling these transitions), rather than a simple rhythm generator.

An important implication of the *state machine* viewpoint is that the RG defines operations in each phase (or state) independent of the exact phase transition mechanisms. During fictive locomotion, these phase transitions and their timing are fully defined by the intrinsic properties of the RG and interactions between them within the spinal cord circuitry. During actual locomotion, these transitions occur under the control of multiple external signals including somatosensory feedback providing information about the states of the controlled limbs and the body (body mechanics). These signals modulate the timing of state (phase) transitions to provide postural stability and adjust locomotion to the animal’s goals and the environment. These signals may delay or advance the onset of the natural (intrinsic) state transitions or enforce the transition when the intrinsic mechanisms are unable to do this.

In this study, we focused on mechanisms that can critically contribute to the swing-to-stance and stance-to-swing phase transitions in RGs controlling each limb. In general, we assume that the timing of these transitions may depend on, and be controlled by, multiple internal (central interactions within and between RGs) and external factors (sensory feedback from the limbs and body or inputs from other brain structures such as the vestibular signals). We used a simplified mathematical description of mouse locomotion and compared the behavior of several model versions to existing experimental data. This approach allowed us to evaluate and suggest the involvement of different mechanisms in phase/state transitions during locomotion and the role of central interactions in the observed locomotor behaviors. Specifically, we suggest that (1) the timing of swing-to-stance transition in each limb at relatively slow locomotor speeds (during walking and trotting gaits without a suspension phase) is controlled by a signal that conveys a postural imbalance, and (2) interlimb coordination, particularly diagonal hind-fore limb synchronization in these gaits, is secured by the corresponding central interactions within the spinal cord.

### 4.2 Locomotor phase/state transitions. Control of swing duration

Despite the ability of spinal CPG circuits to generate the locomotor-like pattern of rhythmic activity in the absence of sensory feedback, this feedback as well as some supraspinal signals were shown to significantly contribute to the control of durations and timing of phase transitions during actual locomotion [9-11, 62-66]. Feedback control of phase transitions was also used in a few earlier models [19, 67-72]. Moreover, the involvement of sensory feedback in locomotor phase transitions expands the frequency range of locomotor oscillations well beyond a very limited range of frequencies generated during fictive locomotion; at high stepping frequencies, the RG/CPG circuits can even lose their ability to intrinsically generate oscillations and operate exclusively as a state machine.

In all versions of our model, the timing of stance-to-swing transitions in the RG controlling liftoff of each limb was defined by two types of sensory feedback from the homonymous limb, carrying information on limb loading and limb extension. The critical role of these feedback signals for stance-to-swing transitions has been explicitly formulated by Ekeberg and Pearson and was successfully implemented in their model for hindlimb cat locomotion [24, 25]. These transition mechanisms have been experimentally supported in multiple studies of cat locomotion. Specifically, the information from limb unloading was mostly provided by force-dependent feedback from ankle extensor muscles, whereas limb extension information comes from the length-dependent feedback from hip flexor muscles (reviewed in [10, 11]).

In contrast to the stance-to-swing transition, the control of the swing-to-stance transition that initiates limb touchdown and defines swing duration is less understood. Studies of mammalian locomotion from mice to humans indicate that locomotor frequency (or, equivalently, the step cycle period) is mostly controlled through changes in the duration of stance with minimal changes in the duration of swing [34-37, 52, 73-76]. This suggests that, although partly modulated by hip/shoulder angles [77], the swing-to-stance transition is mostly defined by central neural mechanisms (neural interactions within and between the spinal RGs) and is much less dependent on the sensory feedback from the limb (at least during locomotion on a flat horizontal surface). Therefore, in the first version of our model (Model 1), we considered locomotion with a constant swing duration in all limbs. However, as we described in the Results (section 3.2.4) this model was rejected because it could not fit the experimental data on swing duration while maintaining a stable locomotion direction (see section 3.2.5 and the corresponding discussion below).

### 4.3 Model 1 lacks a mechanism that stabilizes the movement direction

One of the advantages and hence novelty of our approach to the analysis of locomotion is the consideration of two-dimensional locomotor behaviors with a particular focus on the system’s ability to maintain a constant direction of movement, rather than artificially restricting locomotion to the movements along a straight line only. This has allowed us to account for the dynamics of body displacement in the frontal plane and body orientation, as well as interactions between the two. We could also analyze the effects of these factors on the stability of locomotion, including maintenance the constant movement direction.

As described above, the control of stance duration in Model 1 is based exclusively on the feedback signals from the homonymous limbs and swing has a fixed duration in all limbs. Our analysis has shown that Model 1 can demonstrate stable locomotion only with relatively short swing durations which does not match the experimentally observed locomotor characteristics in mice (Fig. 4A, B1, B2). An attempt to increase swing duration to the physiologically observed values in this model creates an instability resulting from complex interactions between the body displacement in the frontal plane, the angle between the body and the COM trajectory, and the feedback signals involved in phase transitions. This leads to discoordination of left and right limb movements at realistic swing durations, also manifesting as an inability to keep a constant moving direction, resulting in misorientation of the body and an eventual fall (see Fig. 4C and supplemental video).

### 4.4 Control of balance in Models 2 and 3

Models 2 and 3 successfully overcome the limitations of Model 1. In these models, the timing of the swing-to-stance transition (touchdown), and hence the duration of swing, is defined by a signal characterizing a postural instability/loss of balance of the entire body, affecting swing durations of each limb depending on the limb’s state. Most mammals have an early postnatal period during which they learn how to walk stably without falling [78-80]. The results of this learning and the exact feedback mechanism informing the central spinal controller (CPG) on the critical postural imbalance are not known. In Models 2 and 3, we calculate the signal characterizing postural imbalance as a rate of change of the total weight-bearing force (a sum of vertical components of all ground reaction forces) which decreases with postural imbalance. In other words, the loss of balance in Models 2 and 3, indicated by the imbalance signal is considered when the total supporting force is decreasing faster than at a certain rate that represents a threshold for initiation of touchdown and termination of swing. While we do not speculate here how and where this calculation is performed in the central nervous system, the relevant information is potentially available from the limb loading (cutaneous) sensors and can be extracted either within the spinal circuits or in the cerebellum, as well as come from the vestibular system.

Following this concept, we assume that whenever the general imbalance signal exceeds the threshold, the limb currently swinging for the longest time lands and transitions to stance. This selection does not require any high-level processing and may be based on simple adaptive properties of the neurons comprising individual RGs. In other words, the imbalance signal can be seen as a ramping global inhibitory input to the neurons maintaining the swing phase of each limb. However, the effect of this input would depend on their current excitability. If the excitability of each of those neurons gradually reduces during its active (swing) phase, the first neuron to shut down in response to the ramping inhibition will be the one that was active for the longest time. Interestingly, a combination of these two neuronal functions, i.e., maintaining the swing and stance states of the limb and slow adaptation during each locomotor phase, is a prerequisite for endogenous oscillations that can emerge in individual RGs in the absence of any sensory feedback. As mentioned, such oscillations can indeed be induced in both isolated spinal cords and paralyzed animals (see section 4.1), which provides indirect support to our speculations.

We fully realize that the balance-based mechanism and the corresponding imbalance signal controlling limb touchdown and swing duration implemented in our models are based on a certain degree of speculation. Yet, it is clear that mechanisms of the swing-to-stance transitions should operate in coordination with balance control. There are several possible described pathways and mechanisms that can be involved. Crossed [81, 82] and interlimb [83] reflexes from the limbs that are currently in stance can signal a reduction in support and promote a transition from flexion to extension and consequently trigger the touchdown of the limbs that are currently in swing. Balance control can also be mediated by a more complex mechanism involving supraspinal brain structures, such as the motor cortex [84, 85], cerebellum [86, 87], and/or vestibular system [88-91]. It is likely that a combination of mechanisms and pathways is involved in ensuring the appropriate timing of the swing-to-stance transition.

Particularly, quadrupedal animals with vestibular lesions experience balance impairments and exhibit a shorter swing duration and variability in foot placement [92]. It was shown that genetic suppression of vestibular circuits has profound effects on locomotion. Specifically, vestibular-deficient mice could hardly keep the planned trajectory and demonstrated circling episodes during locomotion [89]. In general, abnormal circling can be induced by unilateral lesions of vestibular nuclei in rodents (reviewed in [90, 93]). Interestingly, a similar locomotor instability with random changes of body directions and/or with circling episodes was produced in our Model 1 when increasing swing duration (Fig. 4C). The same locomotor instability would happen in Models 2 and 3 after removal of balance-based control of swing duration and replacing it with the direct control of the swing duration which would effectively transform those models into Model 1. Moreover, as seen in Fig. 4A, the lack of balance control of the swing-to-stance transition (in Model 1) impacts the stability of locomotion, reducing the range of possible swing durations. This generally corresponds to the experimental finding that the contribution of vestibular inputs to the control of locomotion increases when locomotion is slowing down [91].

Incorporating a balance control of the swing-to-stance transition and swing duration in Models 2 and 3 yields stable locomotion within the physiological ranges of locomotor velocities, duty factors, and swing durations in mice (Figs. 8, 10, 11) [29, 35]. These models indeed demonstrate much longer swing durations while maintaining stable locomotion and posture because the incorporated balance control of the swing duration provides extremely robust anti-phase synchronization of left and right limb movements, thus ensuring gait symmetry and preservation of the locomotor direction. We conclude that our simulations implicitly support the possibility that the swing-to-stance transitions and touchdown mechanisms, controlling limb movement in real mouse locomotion, may involve imbalance-related signals functionally similar to those implemented in our models.

Finally, it must be noted that our modeling study was restricted to relatively slow locomotor gaits in which stance durations are not shorter than swing durations (with a duty factors ≥ 0.5). Therefore, the suggested role of the balance-based control of swing can be discussed only in connection with such gaits. During faster gaits (fast trot, galop, or bound), in which animals exhibit suspension phases, load information from the limbs isn’t available at the time of touchdown, and imbalance information cannot be used for the swing-to-stance transition. The role of vestibulospinal control is also thought to be decreased with speed [91]. Thus, at higher-speed gaits, the swing-to-stance transition is likely controlled differently and local somatosensory feedback from each limb sensing the touchdown could be sufficient to maintain stability. For example, cutaneous feedback signals from the plantar surface of the paws (and Ib signals from extensor muscles) can trigger a phase advance of the rhythm generators when stimulated during late swing [94], and a phase-dependent modulation of these cutaneous reflexes results in activation of extensor muscles at the swing-to-stance transition [95, 96].

### 4.5 Limb coordination, locomotor gait changes, and related model limitations

Limb coordination during locomotion, and hence the locomotor gait, can depend on, and be influenced by, both central neural interactions between RG circuits controlling each limb and multiple feedback signals to the RGs reflecting interactions between the neural controller and mechanical components of the system. The latter creates additional interactions between the RGs mediated by peripheral feedback [17, 18, 20, 22, 97, 98]. Since both these interactions depend on the locomotor speed, limb coordination and therefore locomotor gaits are also speed dependent. With an increase in velocity, quadrupeds sequentially switch gaits from walk to trot and then to gallop and bound [45, 99], which has been studied in detail for mice [30, 31]. The role of central (spinal) interactions in these gait transitions has been previously demonstrated in mice [3, 30, 40, 41] and reproduced in a series of computational models [5-7, 37, 39, 40, 53].

The simplified models presented here were limited to walking gaits which we define as gaits without suspension phases. Modeling locomotion at faster (running) gaits, i.e., those involving suspension, such as gallop and bound, requires a 3D description of the mechanics which is a lot more complex. These locomotor behaviors are out of the scope of the present computational study.

Considering the above, our modeling relates to mouse locomotion with duty factors above 0.5, and therefore we excluded gaits that we defined as running [29-31, 35]. In terms of swing duration and speed, this corresponds to the points below the yellow curve representing the duty factor of 0.5 in Fig. 10A. For the points near this curve, e.g., point B2, the locomotor gait is close to ‘pure trot’ with nearly perfect diagonal limb synchronization (Fig. 10B2). At the points on the cyan curve in Fig. 10A (corresponding to a duty factor of 0.75), the locomotor gait is a ‘pure walk’ with persistent 3-leg support during locomotion (Fig. 10B1). Between the yellow and cyan boundaries, the model exhibits a walking gait with alternating 2–leg and 3-leg support phases within the step cycle. With an increase of locomotor speed along the green line in Fig. 10A (or any other line connecting the cyan and yellow curves) the gait continuously transforms from lateral walk to trot (see Fig. 10B1, B2, C2 and the supplemental video [model3.mp4](https://drive.google.com/file/d/13hx0M9_-nZ4EMQUPwU0UpOoUyGuHAeD/view?usp=drive_link)).

One of our goals was to match the main locomotor characteristics, such as stance and swing durations, locomotor frequency, and duty factor, as well as their changes with locomotor speed, to the corresponding experimental data, particularly those published by Herbin et al. [29] and Mendes et al. [35] for overground locomotion, which included locomotor velocities above 20 cm/s. In the model, with the specific linear relationship of the imbalance threshold and the propulsion force (that defines the locomotor velocity) shown in Fig. 10C1, the values of these characteristics and their changes with velocity have good correspondence with the experimental data mentioned (see Fig. 11).

It is important to mention that mice can walk more slowly, i.e., at velocities below 20 cm/s (9±5 cm/s in selected episodes during overground locomotion [30] and 5/10/15 cm/s during treadmill locomotion [31, 100]). It should be noted, however, that such slow walking usually occurs during short episodes of free locomotion or during slow treadmill locomotion. When walking at such a low speed, mice dramatically reduce the stride length to 3.4 ± 0.6 cm during overground locomotion [30], or to less than 4 cm at 5/10/15 cm/s treadmill speeds [31]. Because of the fixed value of the maximal limb displacement used (*D* = 5 cm), our model could produce such slow locomotion at velocities below 20 cm/s with swing duration substantially longer than the one observed in the mentioned experimental studies (0.15-0.2 s vs. ∼0.1 s). Our model can however reproduce walking at such low speeds with realistic swing durations if we reduce the maximal limb displacement to 3 cm, as demonstrated in Fig. 10A. Therefore, we suggest that in order to move with such a low speed (e.g., during exploratory locomotor behavior), mice reduce maximal limb extension which in turn decreases the stride length.

### 4.6 Limb coordination, the role of biomechanics and central neural interactions

Limb coordination during locomotion, and hence the gaits used by animals, depend on many factors such as synaptic drive from supraspinal structures (motor cortex, vestibular nuclei, cerebellum, and brainstem) to spinal circuits [85-87, 89, 91], propriospinal feedback including that from non-homonymous limbs forming crossed and interlimb reflexes [81, 82, 101], central neural interactions between spinal circuits controlling different limbs [3, 30, 40, 41], and mechanical interactions between the body, limbs and the environment [97, 98]. These factors affect limb coordination and produce complex synergistic or antagonistic effects, and their individual contributions depend on other locomotor characteristics such as velocity, phase durations, duty factors, etc.

#### 4.6.1 Locomotion without central neural interactions

In all our models, including Model 1, pure mechanical interactions contribute to left-right anti-phase and diagonal in-phase synchronization of swinging limbs. However, Model 1, which operates without balance-based control of swing duration (unlike Models 2 and 3) and without any central interactions between RGs (unlike Model 3), could only demonstrate a steady symmetric locomotor gait, in which the diagonal limbs are swinging simultaneously. In such a regime, body support is provided by either two diagonal limbs (short periods) or all four limbs (longer part of the step cycle) (see Fig. 4B1, B2). Considering the very large duty factors and short swing durations, such a gait is rather unusual for mouse locomotion [31, 33].

Incorporating the balance-based control of swing duration without central interactions (Model 2) allows locomotion with much longer swing durations and relatively low duty factors (Fig. 8). However, in this model, an increase in swing duration (imbalance threshold), or an increase in velocity, or a reduction of duty factor, transforms the locomotor gait from lateral-sequence walk to pace (Fig. 8B1, B2), which is also unusual for mouse locomotion [30, 31].

Based on the above, realistic locomotor behaviors, including the proper expression of locomotor gaits and their changes, required incorporating specific central interactions between rhythm-generating (RG) circuits controlling each limb as was demonstrated in our Model 3.

#### 4.6.2 The role of central neural interactions

Central interactions between RGs controlling each limb are provided by commissural interneurons (CINs, projecting their axons to the opposite side of the spinal cord) and long descending and ascending propriospinal neurons (LPN, projecting their axons from the cervical to the lumbar enlargement or vice-versa) that mediate interactions between left-right and cervical-lumbar (fore-hind) circuits. Recent molecular genetic studies led to the identification of candidate CINs and LPNs for limb coordination [3, 30, 40, 41, 102-104]. Specifically, the genetically-defined V0 CINs (V0_D_ and V0_V_ types) are involved in left–right alternation in a speed-dependent manner. The ablation of both V0 CIN types (V0_D_ and V0_V_) led to a complete loss of walk, trot, and gallop, leaving bound as the default gait regardless of speed [3, 30, 41]. However, the selective genetic ablation of V0_V_ CINs completely removed the expression of trot, but left intact walk, gallop, and bound. Hence, V0_D_ CINs are essential for left–right alternation at slow locomotor speeds (walk), whereas V0_V_ CINs secure left-right alternation at higher speeds (trot) [3, 41]. Similarly, optogenetic silencing of V3 neurons, including the ascending propriospinal diagonal V3 aLPN (aV3), made the mice unable to move using stable trot, gallop, or bound; these mice could only move with relatively low speed and predominantly used lateral walk [40].

Several computational models were developed to reproduce the above experimental data [5-7, 36-40, 53]. The major central interactions between RGs in the spinal cord proposed in these models, summarized in Fig. 12, include: (a) the left-right excitatory interactions within the cervical and lumbar enlargements mediated by V3 CINs, which are necessary for the expression of asymmetrical gaits (gallop and bound); (b) the left-right inhibitory interactions within the cervical and lumbar enlargements mediated by V0 CINs, which overcome the effects of V3-mediated excitation at low and medium speeds and are necessary for the expression of alternating gaits, such as walk (V0_D_) and trot (V0_V_); and (c) the ascending diagonal V3 aLPNs (aV3) that are necessary for the expression of trot.

**Figure 12.**
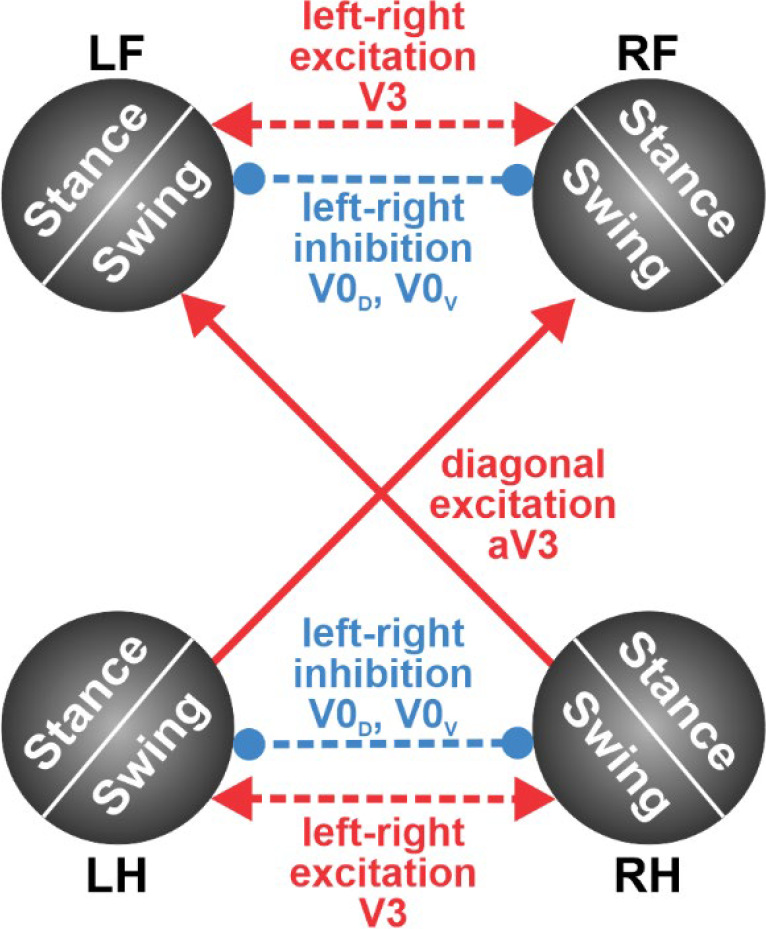
Main central interactions suggested in computational models of spinal locomotor circuits. Red arrows indicate excitatory connections; blue connections with circle ends indicate inhibitory interactions. The dashed lines indicate connections not included in the present model.

Since the present modeling study only focused on locomotion with relatively slow symmetrical gaits with duty factors exceeding 0.5, high-speed gaits such as running trot, gallop, and bound were out of scope. Therefore, the left-right excitatory interactions between circuits controlling homologous RGs, that are likely mediated by V3 CINs, have not been included in the model. For the same reason, we did not include the V0-driven left-right inhibitory interactions between the homologous RGs that should overcome the above excitatory interactions at low velocities. The left-right alternation in all models was only provided by the mechanical interactions. The incorporation of the aV3-mediated ascending diagonal excitation was based on the studies of Zhang et al. [40] (see Fig. 12). Incorporating the corresponding connections has allowed Model 3 to avoid locomotion with a pacing gait not normally observed in mouse locomotion [30, 31]. Despite multiple simplifications, our Model 3, which included diagonal excitation, showed good correspondence to the experimental data (Fig. 11).

### 4.7 Model limitations and future directions

The simplified modeling formalism used here has provided important insights into, and general interpretation of, the neuromechanical control of locomotion in small quadrupedal animals, particularly in mice. Admittedly, our study simulates and analyses mouse locomotion at a restricted range of speeds at which animals usually use symmetrical gaits. While this already covers an important range of natural behaviors, the fast-running trot, gallop, and bound are also frequently expressed in mice, notably during escape, and were not considered in the present study. An extension of the model to these locomotor gaits will require formulating a more complicated description of the movement mechanics. In addition, it will be important for future models to consider additional neural and mechanical factors that could contribute to the control of limb coordination and phase transitions, including reflex circuits, cutaneous feedback, etc. The underlying cell-types and circuits that have been characterized in mice [2] can be added in future models. Finally, some parameters of the model, notably the maximal limb displacement during stance, foot placement locations relative to the body, and body shape, were considered constant. Yet, animals can dynamically change these parameters during overground walking, notably when performing complex and adaptive locomotor maneuvers, such as changing moving trajectory [28], avoiding obstacles, or walking on uneven surfaces. Nevertheless, the modeling framework introduced here can serve as a basis for future modeling and analysis of more complicated and adaptive locomotor behaviors.

## Acknowledgments

This work was supported by NSF CRCNS/DARE grant 2113069 (Ausborn, Rybak), NIH/NINDS grant R01 NS110550 (Rybak), NIH/NINDS grant R01NS112304 (Danner), NIH/NINDS grant R01NS115900 (Danner), CNRS, Agence Nationale de la Recherche (ANR-21-NEUC-0001 and ANR-20-CE16-0026) and Fondation pour la Recherche Médicale (FRM, EQU202203014620) (Bouvier).

## Notes

### Competing Interest Statement

The authors have declared no competing interest.

## References

1. Grillner S. Biological pattern generation: the cellular and computational logic of networks in motion. Neuron. 2006;52(5):751-66. Epub 2006/12/06. doi: S0896-6273(06)00902-0 [pii] 10.1016/j.neuron.2006.11.008. PubMed PMID: 17145498.

2. Kiehn O. Locomotor circuits in the mammalian spinal cord. Annu Rev Neurosci. 2006;29:279-306. Epub 2006/06/17. doi: 10.1146/annurev.neuro.29.051605.112910. PubMed PMID: 16776587.

3. Kiehn O. Decoding the organization of spinal circuits that control locomotion. Nat Rev Neurosci. 2016;17(4):224–38. doi: 10.1038/nrn.2016.9. PubMed PMID: 26935168; PubMed Central PMCID: PMCPMC4844028.

4. Stuart DG, Hultborn H. Thomas Graham Brown (1882–1965), Anders Lundberg (1920–), and the neural control of stepping. Brain research reviews. 2008;59(1):74–95.

5. Rybak IA, Dougherty KJ, Shevtsova NA. Organization of the Mammalian Locomotor CPG: Review of Computational Model and Circuit Architectures Based on Genetically Identified Spinal Interneurons. eNeuro. 2015;2(5):ENEURO.0069-15.2015. doi: 10.1523/ENEURO.0069-15.2015.

6. Danner SM, Wilshin SD, Shevtsova NA, Rybak IA. Central control of interlimb coordination and speed-dependent gait expression in quadrupeds. J Physiol. 2016;594(23):6947–67. doi: 10.1113/JP272787.

7. Danner SM, Shevtsova NA, Frigon A, Rybak IA. Computational modeling of spinal circuits controlling limb coordination and gaits in quadrupeds. eLife. 2017;6:e31050. doi: 10.7554/eLife.31050.

8. Orlovsky GN, Deliagina T, Grillner S. Neuronal control of locomotion: from mollusc to man: Oxford University Press; 1999.

9. Pearson KG. Generating the walking gait: role of sensory feedback. Progress in brain research. 2004;143:123–9. doi: 10.1016/S0079-6123(03)43012-4. PubMed PMID: 14653157.

10. Pearson KG. Role of sensory feedback in the control of stance duration in walking cats. Brain research reviews. 2008;57(1):222-7. Epub 20070729. doi: 10.1016/j.brainresrev.2007.06.014. PubMed PMID: 17761295.

11. Frigon A, Akay T, Prilutsky BI. Control of Mammalian Locomotion by Somatosensory Feedback. Compr Physiol. 2021;12(1):2877-947. Epub 20211229. doi: 10.1002/cphy.c210020. PubMed PMID: 34964114; PubMed Central PMCID: PMCPMC9159344.

12. Kiehn O, Butt SJ. Physiological, anatomical and genetic identification of CPG neurons in the developing mammalian spinal cord. Prog Neurobiol. 2003;70(4):347–61. doi: 10.1016/s0301-0082(03)00091-1. PubMed PMID: 12963092.

13. Juvin L, Simmers J, Morin D. Propriospinal circuitry underlying interlimb coordination in mammalian quadrupedal locomotion. J Neurosci. 2005;25(25):6025–35. doi: 10.1523/JNEUROSCI.0696-05.2005. PubMed PMID: 15976092; PubMed Central PMCID: PMCPMC6724791.

14. Juvin L, Simmers J, Morin D. Locomotor rhythmogenesis in the isolated rat spinal cord: a phase-coupled set of symmetrical flexion extension oscillators. The Journal of physiology. 2007;583(Pt 1):115-28. Epub 20070614. doi: 10.1113/jphysiol.2007.133413. PubMed PMID: 17569737; PubMed Central PMCID: PMCPMC2277226.

15. Cowley KC, Zaporozhets E, Schmidt BJ. Propriospinal transmission of the locomotor command signal in the neonatal rat. Annals of the New York Academy of Sciences. 2010;1198:42–53. doi: 10.1111/j.1749-6632.2009.05421.x. PubMed PMID: 20536919.

16. Zaporozhets E, Cowley KC, Schmidt BJ. Neurochemical excitation of propriospinal neurons facilitates locomotor command signal transmission in the lesioned spinal cord. J Neurophysiol. 2011;105(6):2818-29. Epub 20110330. doi: 10.1152/jn.00917.2010. PubMed PMID: 21451056.

17. Kuo AD. The relative roles of feedforward and feedback in the control of rhythmic movements. Motor Control. 2002;6(2):129–45. doi: 10.1123/mcj.6.2.129. PubMed PMID: 12122223.

18. Righetti L, Ijspeert AJ. Pattern generators with sensory feedback for the control of quadruped locomotion. 2008 IEEE International Conference on Robotics and Automation; 2008 19-23 May 2008.

19. Markin SN, Klishko AN, Shevtsova NA, Lemay MA, Prilutsky BI, Rybak IA. Afferent control of locomotor CPG: insights from a simple neuromechanical model. Annals of the New York Academy of Sciences. 2010;1198:21–34. doi: 10.1111/j.1749-6632.2010.05435.x. PubMed PMID: 20536917.

20. Dzeladini F, van den Kieboom J, Ijspeert A. The contribution of a central pattern generator in a reflex-based neuromuscular model. Front Hum Neurosci. 2014;8(June):1–18. doi: 10.3389/fnhum.2014.00371.

21. Ryu HX, Kuo AD. An optimality principle for locomotor central pattern generators. Sci Rep. 2021;11(1):13140. Epub 20210623. doi: 10.1038/s41598-021-91714-1. PubMed PMID: 34162903; PubMed Central PMCID: PMCPMC8222298.

22. Ijspeert AJ, Daley MA. Integration of feedforward and feedback control in the neuromechanics of vertebrate locomotion: a review of experimental, simulation and robotic studies. J Exp Biol. 2023;226(15). Epub 20230811. doi: 10.1242/jeb.245784. PubMed PMID: 37565347.

23. Di Russo A, Stanev D, Sabnis A, Danner SM, Ausborn J, Armand S, et al. Investigating the roles of reflexes and central pattern generators in the control and modulation of human locomotion using a physiologically plausible neuromechanical model. Journal of neural engineering. 2023. Epub 20230927. doi: 10.1088/1741-2552/acfdcc. PubMed PMID: 37757805.

24. Ekeberg O, Pearson K. Computer simulation of stepping in the hind legs of the cat: an examination of mechanisms regulating the stance-to-swing transition. J Neurophysiol. 2005;94(6):4256-68. Epub 20050727. doi: 10.1152/jn.00065.2005. PubMed PMID: 16049149.

25. Pearson K, Ekeberg O, Buschges A. Assessing sensory function in locomotor systems using neuro-mechanical simulations. Trends Neurosci. 2006;29(11):625-31. Epub 20060907. doi: 10.1016/j.tins.2006.08.007. PubMed PMID: 16956675.

26. Geyer H, Herr H. A Muscle-Reflex Model That Encodes Principles of Legged Mechanics Produces Human Walking Dynamics and Muscle Activities. IEEE Transactions on Neural Systems and Rehabilitation Engineering. 2010;18(3):263–73. doi: 10.1109/TNSRE.2010.2047592.

27. Song S, Geyer H. A neural circuitry that emphasizes spinal feedback generates diverse behaviours of human locomotion. The Journal of physiology. 2015;593(16):3493-511. Epub 20150623. doi: 10.1113/JP270228. PubMed PMID: 25920414; PubMed Central PMCID: PMCPMC4560581.

28. Gruntman E, Benjamini Y, Golani I. Coordination of steering in a free-trotting quadruped. J Comp Physiol A Neuroethol Sens Neural Behav Physiol. 2007;193(3):331-45. Epub 20061205. doi: 10.1007/s00359-006-0187-5. PubMed PMID: 17146663.

29. Herbin M, Hackert R, Gasc JP, Renous S. Gait parameters of treadmill versus overground locomotion in mouse. Behav Brain Res. 2007;181(2):173-9. Epub 20070408. doi: 10.1016/j.bbr.2007.04.001. PubMed PMID: 17521749.

30. Bellardita C, Kiehn O. Phenotypic characterization of speed-associated gait changes in mice reveals modular organization of locomotor networks. Curr Biol. 2015;25(11):1426-36. Epub 20150507. doi: 10.1016/j.cub.2015.04.005. PubMed PMID: 25959968; PubMed Central PMCID: PMCPMC4469368.

31. Lemieux M, Josset N, Roussel M, Couraud S, Bretzner F. Speed-Dependent Modulation of the Locomotor Behavior in Adult Mice Reveals Attractor and Transitional Gaits. Front Neurosci. 2016;10:42. Epub 20160223. doi: 10.3389/fnins.2016.00042. PubMed PMID: 26941592; PubMed Central PMCID: PMCPMC4763020.

32. Ekeberg O, Pearson K. Computer simulation of stepping in the hind legs of the cat: an examination of mechanisms regulating the stance-to-swing transition. J Neurophysiol. 2005;94(6):4256–68. doi: 10.1152/jn.00065.2005.

33. Garrick JM, Costa LG, Cole TB, Marsillach J. Evaluating Gait and Locomotion in Rodents with the CatWalk. Current Protocols. 2021;1(8):e220. doi: 10.1002/cpz1.220.

34. Batka RJ, Brown TJ, McMillan KP, Meadows RM, Jones KJ, Haulcomb MM. The need for speed in rodent locomotion analyses. Anatomical record. 2014;297(10):1839-64. Epub 20140603. doi: 10.1002/ar.22955. PubMed PMID: 24890845; PubMed Central PMCID: PMCPMC4758221.

35. Mendes CS, Bartos I, Marka Z, Akay T, Marka S, Mann RS. Quantification of gait parameters in freely walking rodents. BMC Biol. 2015;13:50. Epub 20150722. doi: 10.1186/s12915-015-0154-0. PubMed PMID: 26197889; PubMed Central PMCID: PMCPMC4511453.

36. Molkov YI, Bacak BJ, Talpalar AE, Rybak IA. Mechanisms of left-right coordination in mammalian locomotor pattern generation circuits: a mathematical modeling view. PLoS Comput Biol. 2015;11(5):e1004270. doi: 10.1371/journal.pcbi.1004270. PubMed PMID: 25970489; PubMed Central PMCID: PMCPMC4430237.

37. Shevtsova NA, Talpalar AE, Markin SN, Harris-Warrick RM, Kiehn O, Rybak IA. Organization of left-right coordination of neuronal activity in the mammalian spinal cord: Insights from computational modelling. The Journal of physiology. 2015;593(11):2403-26. Epub 2015/03/31. doi: 10.1113/JP270121. PubMed PMID: 25820677; PubMed Central PMCID: PMCPMC4461406.

38. Ausborn J, Shevtsova NA, Caggiano V, Danner SM, Rybak IA. Computational modeling of brainstem circuits controlling locomotor frequency and gait. Elife. 2019;8. Epub 2019/01/22. doi: 10.7554/eLife.43587. PubMed PMID: 30663578; PubMed Central PMCID: PMCPMC6355193.

39. Danner SM, Zhang H, Shevtsova NA, Borowska-Fielding J, Deska-Gauthier D, Rybak IA, et al. Spinal V3 Interneurons and Left-Right Coordination in Mammalian Locomotion. Front Cell Neurosci. 2019;13:516. Epub 2019/12/12. doi: 10.3389/fncel.2019.00516. PubMed PMID: 31824266; PubMed Central PMCID: PMCPMC6879559.

40. Zhang H, Shevtsova NA, Deska-Gauthier D, Mackay C, Dougherty KJ, Danner SM, et al. The role of V3 neurons in speed-dependent interlimb coordination during locomotion in mice. Elife. 2022;11. Epub 20220427. doi: 10.7554/eLife.73424. PubMed PMID: 35476640; PubMed Central PMCID: PMCPMC9045817.

41. Talpalar AE, Bouvier J, Borgius L, Fortin G, Pierani A, Kiehn O. Dual-mode operation of neuronal networks involved in left-right alternation. Nature. 2013;500(7460):85–8. doi: 10.1038/nature12286. PubMed PMID: 23812590.

42. Bellardita C, Kiehn O. Phenotypic Characterization of Speed-Associated Gait Changes in Mice Reveals Modular Organization of Locomotor Networks. Curr Biol. 2015;25(11):1426–36. doi: 10.1016/j.cub.2015.04.005.

43. Ruder L, Takeoka A, Arber S. Long-Distance Descending Spinal Neurons Ensure Quadrupedal Locomotor Stability. Neuron. 2016;92(5):1063–78. doi: 10.1016/j.neuron.2016.10.032.

44. Alexander RM. Mechanics and scaling of terrestrial locomotion. In: Pedley TJ, editor. Scale effects in animal locomotion. London: Academic Press; 1977. p. 93-110.

45. Hildebrand M. The Quadrupedal Gaits of Vertebrates: The timing of leg movements relates to balance, body shape, agility, speed, and energy expenditure. BioScience. 1989;39(11):766–75. doi: 10.2307/1311182.

46. Frigon A. The neural control of interlimb coordination during mammalian locomotion. J Neurophysiol. 2017;117(6):2224-41. Epub 2017/03/17. doi: 10.1152/jn.00978.2016. PubMed PMID: 28298308; PubMed Central PMCID: PMCPMC5454475.

47. Brown TG. On the nature of the fundamental activity of the nervous centres; together with an analysis of the conditioning of rhythmic activity in progression, and a theory of the evolution of function in the nervous system. The Journal of physiology. 1914;48(1):18-46. Epub 1914/03/31. doi: 10.1113/jphysiol.1914.sp001646. PubMed PMID: 16993247; PubMed Central PMCID: PMCPMC1420503.

48. Grillner S, Brookhart M, Mountcastle V. Handbook of Physiology, section 1, The Nervous System, vol. II, Motor Control. American Physiological Society; 1981.

49. Hultborn H, Nielsen JB. Spinal control of locomotion--from cat to man. Acta Physiol (Oxf). 2007;189(2):111–21. doi: 10.1111/j.1748-1716.2006.01651.x. PubMed PMID: 17250563.

50. Lundberg A. HALF-CENTRES REVISITED. In: Szentágothai J, Palkovits M, Hámori J, editors. Regulatory Functions of the CNS Principles of Motion and Organization: Pergamon; 1981. p. 155–67.

51. Rossignol S. Visuomotor regulation of locomotion. Canadian journal of physiology and pharmacology. 1996;74(4):418–25.

52. McCrea DA, Rybak IA. Organization of mammalian locomotor rhythm and pattern generation. Brain Res Rev. 2008;57(1):134–46. doi: 10.1016/j.brainresrev.2007.08.006.

53. Ausborn J, Shevtsova NA, Danner SM. Computational Modeling of Spinal Locomotor Circuitry in the Age of Molecular Genetics. Int J Mol Sci. 2021;22(13):6835. PubMed PMID: doi:10.3390/ijms22136835.

54. Grillner S, Zangger P. On the central generation of locomotion in the low spinal cat. Exp Brain Res. 1979;34(2):241–61. doi: 10.1007/BF00235671. PubMed PMID: 421750.

55. McCrea DA. Spinal circuitry of sensorimotor control of locomotion. The Journal of physiology. 2001;533(Pt 1):41–50. doi: 10.1111/j.1469-7793.2001.0041b.x. PubMed PMID: 11351011; PubMed Central PMCID: PMCPMC2278617.

56. Lafreniere-Roula M, McCrea DA. Deletions of rhythmic motoneuron activity during fictive locomotion and scratch provide clues to the organization of the mammalian central pattern generator. J Neurophysiol. 2005;94(2):1120-32. Epub 20050504. doi: 10.1152/jn.00216.2005. PubMed PMID: 15872066.

57. Smith JC, Feldman JL, Schmidt BJ. Neural mechanisms generating locomotion studied in mammalian brain stem-spinal cord in vitro. Faseb J. 1988;2(7):2283–8. doi: 10.1096/fasebj.2.7.2450802. PubMed PMID: 2450802.

58. Cowley KC, Schmidt BJ. Regional distribution of the locomotor pattern-generating network in the neonatal rat spinal cord. J Neurophysiol. 1997;77(1):247-59. Epub 1997/01/01. doi: 10.1152/jn.1997.77.1.247. PubMed PMID: 9120567.

59. Ding R, Yu J, Yang Q, Tan M, Zhang J. CPG-based behavior design and implementation for a biomimetic amphibious robot. 2011 IEEE International Conference on Robotics and Automation; 2011 9-13 May 2011.

60. Liang D, Kreiser R, Nielsen C, Qiao N, Sandamirskaya Y, Indiveri G. Neural State Machines for Robust Learning and Control of Neuromorphic Agents. IEEE Journal on Emerging and Selected Topics in Circuits and Systems. 2019;9(4):679–89. doi: 10.1109/JETCAS.2019.2951442.

61. Spaeth A, Tebyani M, Haussler D, Teodorescu M. Spiking neural state machine for gait frequency entrainment in a flexible modular robot. PloS one. 2020;15(10):e0240267. Epub 20201021. doi: 10.1371/journal.pone.0240267. PubMed PMID: 33085673; PubMed Central PMCID: PMCPMC7577446.

62. Grillner S, Rossignol S. On the initiation of the swing phase of locomotion in chronic spinal cats. Brain Res. 1978;146(2):269–77. doi: 10.1016/0006-8993(78)90973-3. PubMed PMID: 274169.

63. Hiebert GW, Whelan PJ, Prochazka A, Pearson KG. Contribution of hind limb flexor muscle afferents to the timing of phase transitions in the cat step cycle. J Neurophysiol. 1996;75(3):1126–37. doi: 10.1152/jn.1996.75.3.1126. PubMed PMID: 8867123.

64. Duysens J, Pearson KG. Inhibition of flexor burst generation by loading ankle extensor muscles in walking cats. Brain Res. 1980;187(2):321–32. doi: 10.1016/0006-8993(80)90206-1. PubMed PMID: 7370733.

65. Lam T, Pearson KG. Proprioceptive modulation of hip flexor activity during the swing phase of locomotion in decerebrate cats. J Neurophysiol. 2001;86(3):1321–32. doi: 10.1152/jn.2001.86.3.1321. PubMed PMID: 11535680.

66. Lam T, Pearson KG. Sartorius muscle afferents influence the amplitude and timing of flexor activity in walking decerebrate cats. Exp Brain Res. 2002;147(2):175-85. Epub 20020913. doi: 10.1007/s00221-002-1236-0. PubMed PMID: 12410332.

67. Taga G. A model of the neuro-musculo-skeletal system for anticipatory adjustment of human locomotion during obstacle avoidance. Biol Cybern. 1998;78(1):9–17. doi: 10.1007/s004220050408. PubMed PMID: 9485584.

68. Ivashko DG, Prilutsky BI, Markin S, Chapin JK, Rybak IA. Modeling the spinal cord neural circuitry controlling cat hindlimb movement during locomotion. Neurocomputing. 2003;52:621–9.

69. Yakovenko S, Gritsenko V, Prochazka A. Contribution of stretch reflexes to locomotor control: a modeling study. Biol Cybern. 2004;90(2):146-55. Epub 20040120. doi: 10.1007/s00422-003-0449-z. PubMed PMID: 14999481.

70. Markin SN, Klishko AN, Shevtsova NA, Lemay MA, Prilutsky BI, Rybak IA. A Neuromechanical Model of Spinal Control of Locomotion. In: Prilutsky BI, Edwards DH, editors. Neuromechanical Model Posture Locomot. New York, NY: Springer New York; 2016. p. 21–65.

71. Aoi S, Ogihara N, Funato T, Sugimoto Y, Tsuchiya K. Evaluating functional roles of phase resetting in generation of adaptive human bipedal walking with a physiologically based model of the spinal pattern generator. Biol Cybern. 2010;102(5):373-87. Epub 20100309. doi: 10.1007/s00422-010-0373-y. PubMed PMID: 20217427.

72. Aoi S, Kondo T, Hayashi N, Yanagihara D, Aoki S, Yamaura H, et al. Contributions of phase resetting and interlimb coordination to the adaptive control of hindlimb obstacle avoidance during locomotion in rats: a simulation study. Biol Cybern. 2013;107(2):201–16. doi: 10.1007/s00422-013-0546-6. PubMed PMID: 23430278.

73. Halbertsma JM. The stride cycle of the cat: the modelling of locomotion by computerized analysis of automatic recordings. Acta physiologica Scandinavica Supplementum. 1983;521:1-75. PubMed PMID: 6582764.

74. Leblond H, L’Esperance M, Orsal D, Rossignol S. Treadmill locomotion in the intact and spinal mouse. J Neurosci. 2003;23(36):11411–9. doi: 10.1523/JNEUROSCI.23-36-11411.2003. PubMed PMID: 14673005; PubMed Central PMCID: PMCPMC6740531.

75. Frigon A, Gossard JP. Asymmetric control of cycle period by the spinal locomotor rhythm generator in the adult cat. The Journal of physiology. 2009;587(Pt 19):4617-28. Epub 2009/08/14. doi: 10.1113/jphysiol.2009.176669. PubMed PMID: 19675066; PubMed Central PMCID: PMCPMC2768017.

76. Frigon A, Hurteau M-F, Thibaudier Y, Leblond H, Telonio A, D’Angelo G. Split-belt walking alters the relationship between locomotor phases and cycle duration across speeds in intact and chronic spinalized adult cats. Journal of Neuroscience. 2013;33(19):8559–66.

77. McVea DA, Donelan JM, Tachibana A, Pearson KG. A role for hip position in initiating the swing-to-stance transition in walking cats. J Neurophysiol. 2005;94(5):3497-508. Epub 20050810. doi: 10.1152/jn.00511.2005. PubMed PMID: 16093331.

78. Garwicz M, Christensson M, Psouni E. A unifying model for timing of walking onset in humans and other mammals. Proceedings of the National Academy of Sciences. 2009;106(51):21889–93. doi: 10.1073/pnas.0905777106.

79. Shriner AM, Drever FR, Metz GA. The development of skilled walking in the rat. Behav Brain Res. 2009;205(2):426-35. Epub 20090804. doi: 10.1016/j.bbr.2009.07.029. PubMed PMID: 19660502; PubMed Central PMCID: PMCPMC5222626.

80. Nguyen KP, Sharma A, Gil-Silva M, Gittis AH, Chase SM. Distinct Kinematic Adjustments over Multiple Timescales Accompany Locomotor Skill Development in Mice. Neuroscience. 2021;466:260-72. Epub 20210524. doi: 10.1016/j.neuroscience.2021.05.002. PubMed PMID: 34088581; PubMed Central PMCID: PMCPMC8561674.

81. Laflamme OD, Ibrahim M, Akay T. Crossed reflex responses to flexor nerve stimulation in mice. J Neurophysiol. 2022;127(2):493-503. Epub 20220105. doi: 10.1152/jn.00385.2021. PubMed PMID: 34986055; PubMed Central PMCID: PMCPMC8836714.

82. Laflamme OD, Markin SN, Deska-Gauthier D, Banks R, Zhang Y, Danner SM, et al. Distinct roles of spinal commissural interneurons in transmission of contralateral sensory information. Curr Biol. 2023;33(16):3452-64 e4. Epub 20230801. doi: 10.1016/j.cub.2023.07.014. PubMed PMID: 37531957.

83. Hurteau MF, Frigon A. A Spinal Mechanism Related to Left-Right Symmetry Reduces Cutaneous Reflex Modulation Independently of Speed During Split-Belt Locomotion. J Neurosci. 2018;38(48):10314-28. Epub 2018/10/14. doi: 10.1523/JNEUROSCI.1082-18.2018. PubMed PMID: 30315129.

84. Klishko AN, Farrell BJ, Beloozerova IN, Latash ML, Prilutsky BI. Stabilization of cat paw trajectory during locomotion. J Neurophysiol. 2014;112(6):1376-91. Epub 20140603. doi: 10.1152/jn.00663.2013. PubMed PMID: 24899676; PubMed Central PMCID: PMCPMC4137248.

85. Farrell BJ, Bulgakova MA, Sirota MG, Prilutsky BI, Beloozerova IN. Accurate stepping on a narrow path: mechanics, EMG, and motor cortex activity in the cat. J Neurophysiol. 2015;114(5):2682-702. Epub 2015/09/12. doi: 10.1152/jn.00510.2014. PubMed PMID: 26354314; PubMed Central PMCID: PMCPMC4644224.

86. Machado AS, Darmohray DM, Fayad J, Marques HG, Carey MR. A quantitative framework for whole-body coordination reveals specific deficits in freely walking ataxic mice. Elife. 2015;4. doi: 10.7554/eLife.07892. PubMed PMID: 26433022; PubMed Central PMCID: PMCPMC4630674.

87. Machado AS, Marques HG, Duarte DF, Darmohray DM, Carey MR. Shared and specific signatures of locomotor ataxia in mutant mice. Elife. 2020;9. Epub 20200728. doi: 10.7554/eLife.55356. PubMed PMID: 32718435; PubMed Central PMCID: PMCPMC7386913.

88. Vidal PP, Degallaix L, Josset P, Gasc JP, Cullen KE. Postural and locomotor control in normal and vestibularly deficient mice. The Journal of physiology. 2004;559(Pt 2):625-38. Epub 20040708. doi: 10.1113/jphysiol.2004.063883. PubMed PMID: 15243133; PubMed Central PMCID: PMCPMC1665125.

89. Eugene D, Deforges S, Vibert N, Vidal PP. Vestibular critical period, maturation of central vestibular neurons, and locomotor control. Annals of the New York Academy of Sciences. 2009;1164:180–7. doi: 10.1111/j.1749-6632.2008.03727.x. PubMed PMID: 19645897.

90. Loscher W. Abnormal circling behavior in rat mutants and its relevance to model specific brain dysfunctions. Neurosci Biobehav Rev. 2010;34(1):31-49. Epub 20090714. doi: 10.1016/j.neubiorev.2009.07.001. PubMed PMID: 19607857.

91. Akay T, Murray AJ. Relative Contribution of Proprioceptive and Vestibular Sensory Systems to Locomotion: Opportunities for Discovery in the Age of Molecular Science. Int J Mol Sci. 2021;22(3). Epub 20210202. doi: 10.3390/ijms22031467. PubMed PMID: 33540567; PubMed Central PMCID: PMCPMC7867206.

92. McCrum C, Lucieer F, van de Berg R, Willems P, Perez Fornos A, Guinand N, et al. The walking speed-dependency of gait variability in bilateral vestibulopathy and its association with clinical tests of vestibular function. Sci Rep. 2019;9(1):18392. Epub 20191205. doi: 10.1038/s41598-019-54605-0. PubMed PMID: 31804514; PubMed Central PMCID: PMCPMC6895118.

93. Rastoldo G, Marouane E, El Mahmoudi N, Pericat D, Bourdet A, Timon-David E, et al. Quantitative Evaluation of a New Posturo-Locomotor Phenotype in a Rodent Model of Acute Unilateral Vestibulopathy. Front Neurol. 2020;11:505. Epub 20200605. doi: 10.3389/fneur.2020.00505. PubMed PMID: 32582016; PubMed Central PMCID: PMCPMC7291375.

94. Duysens J. Reflex control of locomotion as revealed by stimulation of cutaneous afferents in spontaneously walking premammillary cats. Journal of neurophysiology. 1977;40(4):737--51. PubMed PMID: Duysens1977.

95. Yang JF, Stein RB. Phase-dependent reflex reversal in human leg muscles during walking. J Neurophysiol. 1990;63(5):1109–17. doi: 10.1152/jn.1990.63.5.1109. PubMed PMID: 2358865.

96. Rossignol S, Dubuc R, Gossard J-P. Dynamic sensorimotor interactions in locomotion. Physiol Rev. 2006;86(1):89–154. doi: 10.1152/physrev.00028.2005.

97. Full RJ, Koditschek DE. Templates and anchors: neuromechanical hypotheses of legged locomotion on land. J Exp Biol. 1999;202(Pt 23):3325-32. doi: 10.1242/jeb.202.23.3325. PubMed PMID: 10562515.

98. Full RJ, Kubow T, Schmitt J, Holmes P, Koditschek D. Quantifying dynamic stability and maneuverability in legged locomotion. Integrative and comparative biology. 2002;42(1):149–57. doi: 10.1093/icb/42.1.149. PubMed PMID: 21708704.

99. Hildebrand M. The Adaptive Significance of Tetrapod Gait Selection. American Zoologist. 1980;20(1):255–67. doi: 10.1093/icb/20.1.255.

100. Herent C, Diem S, Fortin G, Bouvier J. Absent phasing of respiratory and locomotor rhythms in running mice. Elife. 2020;9. Epub 20201201. doi: 10.7554/eLife.61919. PubMed PMID: 33258770; PubMed Central PMCID: PMCPMC7707822.

101. Hurteau MF, Thibaudier Y, Dambreville C, Danner SM, Rybak IA, Frigon A. Intralimb and Interlimb Cutaneous Reflexes during Locomotion in the Intact Cat. J Neurosci. 2018;38(17):4104-22. Epub 2018/03/23. doi: 10.1523/JNEUROSCI.3288-17.2018. PubMed PMID: 29563181; PubMed Central PMCID: PMCPMC5963849.

102. Lanuza GM, Gosgnach S, Pierani A, Jessell TM, Goulding M. Genetic identification of spinal interneurons that coordinate left-right locomotor activity necessary for walking movements. Neuron. 2004;42(3):375–86. doi: 10.1016/s0896-6273(04)00249-1. PubMed PMID: 15134635.

103. Zhang Y, Narayan S, Geiman E, Lanuza GM, Velasquez T, Shanks B, et al. V3 spinal neurons establish a robust and balanced locomotor rhythm during walking. Neuron. 2008;60(1):84–96. doi: 10.1016/j.neuron.2008.09.027. PubMed PMID: 18940590; PubMed Central PMCID: PMCPMC2753604.

104. Goulding M. Circuits controlling vertebrate locomotion: moving in a new direction. Nat Rev Neurosci. 2009;10(7):507–18. doi: 10.1038/nrn2608. PubMed PMID: 19543221; PubMed Central PMCID: PMCPMC2847453.

